# Human enteric viruses shape disease phenotype through divergent innate immunomodulation

**DOI:** 10.1101/2021.10.14.464404

**Authors:** Fatemeh Adiliaghdam, Hajera Amatullah, Sreehaas Digumarthi, Tahnee L. Saunders, Raza-Ur Rahman, Lai Ping Wong, Ruslan Sadreyev, Lindsay Droit, Jean Paquette, Philippe Goyette, John Rioux, Richard Hodin, Kathie A. Mihindukulasuriya, Scot A. Handley, Kate L. Jeffrey

## Abstract

Altered enteric microorganisms in concert with host genetics shape inflammatory bowel disease (IBD) phenotypes. However, insight is limited to bacteria and fungi. We found virus like particles (VLPs) enriched from normal human colon resections, containing eukaryotic viruses and bacteriophages (collectively, the virome), actively elicited atypical anti-inflammatory innate immune programs. Conversely, IBD patient VLPs provoked inflammation, which was successfully dampened by healthy VLPs. The IBD colon tissue virome was perturbed, including enriched Picornovirus *Enterovirus B,* not previously observed in fecal virome studies. Mice with humanized healthy colon tissue viromes had attenuated intestinal inflammation while those with IBD-derived viromes exhibited exacerbated inflammation in a nucleic acid sensing-dependent fashion. Furthermore, there were detrimental consequences for IBD-associated MDA5 loss-of-function on patient intestinal epithelial cells exposed to healthy or IBD viromes. Our results demonstrate that innate recognition of either healthy or IBD human viromes autonomously influences disease phenotypes in IBD. Harnessing the virome may offer therapeutic and biomarker potential.

**One Sentence Summary:** Human viromes divergently shape host immunity and disease

## Main Text

Viruses are the most abundant biological entity on earth and infect all types of life. Generally considered obligate pathogens, viruses are a constant and emerging threat to human health, as exemplified by the Severe Acute Respiratory Syndrome Coronavirus 2 (SARS-CoV-2) pandemic. Yet, paradoxically, many viruses persistently reside within asymptomatic individuals. These include a range of eukaryotic viruses, as well as bacteriophages (*1–4*), that exist both within their bacterial host, as well as free virions embedded within mucus layers of the intestine (*5*). These collective resident viruses, along with bacteria, archaea, fungi, helminths and protozoa form the human metagenome that have intimate functional relationships with their host. The human intestinal virome is established at birth and is dominated by bacteriophages, while eukaryotic viruses gradually emerge after birth, as measured in feces that contains ∼10^9^ virus like particles (VLPs) per gram (*2, 6*). Furthermore, despite multiple remaining hurdles in identifying what constitutes a normal human virome (*7*), robust fluctuations of the virome have been reported in IBD (*1, 8–10*), colorectal cancer (*11, 12*), type I diabetes (*13*), nonalcoholic fatty liver disease (*14*), cystic fibrosis (*15*), graft-versus-host-disease (*16*) and HIV infection (*17*) and a range of other diseases (*18*). However, this rapid cataloging of viral genomes in human tissues by sequencing has vastly outpaced any mechanistic understanding we have of virome biology. Viruses inhabiting a healthy host (*19, 20*), and host innate sensing of viruses by gut-resident Toll-like Receptors (TLRs) or RIG-I like Receptors (RLRs) (*19, 21–26*) provide homeostasis in the mouse intestine. However, functional studies of the human healthy or diseased enteric virome are currently lacking. Although, examples of direct and distinct bacteriophage-human immune interactions are emerging (*5, 27, 28*). Furthermore, loss-of-function variants in the gene *IFIH1* that encodes the host virus receptor melanoma differentiation associated protein 5 (MDA5) were shown to significantly associate with IBD by genome wide-association studies, GWAS (*29*). However, how abrogation of MDA5 function may contribute to intestinal inflammation in IBD, especially in the context of the virome, is unclear.

To determine whether the human enteric virome has homeostatic or pathophysiological properties, we purified virus-like particles (VLPs) from fresh colon resections taken from non-inflamed/non-IBD, Ulcerative Colitis (UC) or Crohn’s disease (CD) patients, post-surgery (**Fig 1A**, **Table S1**). We reasoned that since viruses are obligate intracellular pathogens that rely on a host organism for successful replication, and bacteriophages reside within mucosal layers (*5*) and are readily taken up by eukaryotic cells (*27, 30, 31*) certain viruses may be better represented in upper intestinal tissue than in feces. Tissue from non-IBD individuals was confirmed to be not inflamed and UC and CD tissue was confirmed to be inflamed by a blinded clinical pathologist (**Table S1**). We quantified our VLP preparations by confocal microscopy using nucleic acid-binding SYBR Gold, and found an average of 2.8×10^7^ VLPs per mg of colon tissue from non-IBD, UC or CD individuals (**Fig 1B**). Our preparations were comprised of prokaryotic and eukaryotic viruses, as visualized by electron microscopy (**Fig S1A**), and importantly, human colon-derived VLP preparations were both endotoxin- and cytokine-free (**Fig S1B-F**).

**Fig 1.**
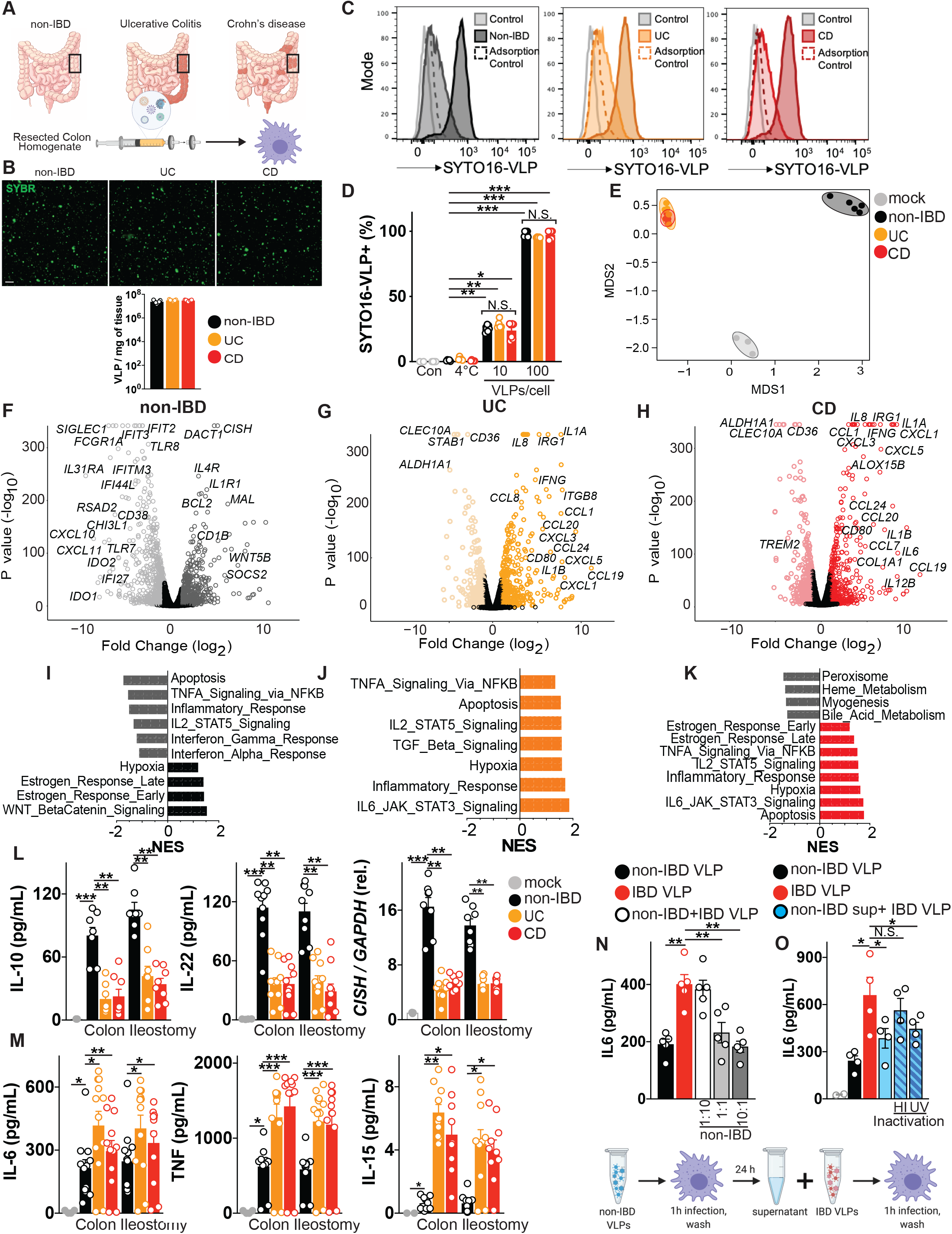
Human colon tissue resident viruses protect while IBD viromes divergently provoke inflammation. **(A)** Virus-like particles (VLPs) isolated from colon resections post-surgery from non-IBD (n=5), Ulcerative Colitis (UC, n=5) or Crohn’s disease (CD, n=5) patients. **(B)** Visualization and quantification of VLP isolates per mg of colon tissue using confocal microscopy for RNA/DNA using SYBR Gold. Scale bar is 1μm. **(C, D)** Flow cytometric quantification of uptake of SYTO-16 labeled human colon-resection derived VLPs by human peripheral blood-derived macrophages (1 or 10 VLPs per cell). Adsorption control is fluorescently labeled VLPs delivered to human macrophages at 4°C. **(E)** Multidimensional scaling (MDS) plot of human macrophage transcriptional profiles induced by non-IBD or UC or CD derived VLPs (10 VLPs/cell, 24h). **(F-H)** Volcano plots showing differentially expressed genes comparing **F)** non-IBD delivery to mock adsorption, **(G)** UC delivery to mock adsorption or **(H)** CD delivery to mock adsorption. **(I-K)** Hallmark gene sets significantly (P value< 0.05, FDR <0.20) represented in **(I)** non-IBD, **(J)** UC or, **(K)** CD VLP induced transcriptional programs as assessed by Gene Set Enrichment Analysis (GSEA). NES is normalized enrichment score. **(L)** Production of anti-inflammatory mediators Interleukin (IL)-10, IL-22 or *CISH* or **(M)** pro-inflammatory mediators TNF, IL-6, IL-15 as measured by ELISA or qPCR following delivery of VLPs (10 VLPs/cell, 24h) extracted from non-IBD, Ulcerative Colitis (UC) or Crohn’s disease (CD) colon resections (n=8 per group) or ileostomy content (n=8 per group) to primary human peripheral blood-derived macrophages. **(N)** Primary human peripheral blood-derived macrophages were delivered colon resection derived non-IBD VLPs (10 VLPs/cell), IBD VLPs (10 VLPs/cell) or indicated ratios of non IBD to IBD VLPs (total of 10 VLPs/cell) for 24 hours and IL-6 was measured by ELISA. (**O**) IL-6 levels measured by ELISA following IBD VLP delivery to human macrophages in the presence of non-IBD macrophage supernatant that had been heat inactivated (95°C 10 min) or UV crosslinked (3x 200mJ/cm^2^) as controls. Data are mean ± s.e.m. of 8-12 biological replicates \**P*<0.05, \*\**P*<0.01, \*\*\**P*<0.001, \*\*\*\**P*<0.0001 as determined by one-way ANOVA with Tukey’s multiple comparison test.

Our understanding of the host innate immune response to virus stems predominantly from studies of resolving acute infection. This mammalian antiviral paradigm centers on the detection of virus moieties by cytosolic or endosomal receptors and production of type I or III interferon (IFNs) and hundreds of IFN stimulated genes (ISGs) that aim to eliminate virus, or the cell that it infects (*32*). The rules governing if and how innate immune cells chronically respond to persistent eukaryotic or prokaryotic viruses in the intestine, as well as in other systemic niches (*33*), and how this shapes host physiology, are still poorly understood. Macrophages are one of the most abundant leukocytes in the intestinal mucosa where they are essential for maintaining homeostasis but are also implicated in the pathogenesis of IBD (*34*). Macrophages are also critical sensors of virus, including endocytosed bacteriophage (*27*). Therefore, to identify the precise innate immune response initiated by eukaryotic or prokaryotic viruses residing in healthy or diseased human colon, we exposed primary human peripheral blood derived macrophages to collective VLPs isolated from non-IBD, UC or CD patient colon resections (**Fig 1A**). Fluorescently labeled colon resection-derived VLPs demonstrated dose-dependent internalization by primary human macrophages that was equal for non-IBD, UC or CD-derived viromes (**Fig 1C, D**). Furthermore, virome uptake was dependent on endocytosis as treatment with brefeldin A a vesicular transport inhibitor, greatly reduced intracellular accumulation (**Fig. S2**). Thus, the human virome, that includes both eukaryotic viruses and bacteriophages, is readily internalized by human immune cells via endocytosis for pathogen associated molecular pattern (PAMP) recognition, consistent with other reports (*27, 30*). To assess the transcriptional pathways induced by the human colon tissue virome in macrophages, we performed unbiased RNA sequencing. Multi-dimensional scaling (MDS) analysis found non-IBD colon-resident viromes initiated a distinct transcriptional program, that was highly reproducible between 5 different patient VLP sources (**Fig 1E**). Conversely, UC or CD human enteric viromes from multiple patients induced macrophage transcriptional programs that were distinct from non-IBD virome-induced programs but almost indistinguishable from one another (**Fig 1E**). Interestingly, viruses resident to normal colon tissue predominantly suppressed the host innate immune response as more transcripts were down-regulated in macrophages exposed to non-IBD virus than were up-regulated (**Fig S3**). On the other hand, both UC and CD viromes triggered a prominent upregulation in gene expression of macrophages and these transcriptional programs were almost identical (**Fig S3**). A striking feature of the non-IBD virome induced macrophage transcriptional signature was a broad downregulation of the classical antiviral response. Non-IBD viromes significantly downregulated an array of virus receptors, signalling adapters and IFN stimuated genes (ISGs) such as *TLR7*, *TLR8*, *IFIT2*, *IFIT3*, *RSAD2* (also known as Viperin) and *IFI44L* (**Fig 1F and Fig S3D**). This atypical host response to non-IBD viromes was also exemplified by the prominent induction of anti-inflammatory and pro-survival genes such as *IL4R*, *CISH*, *BCL2*, *SOCS2* (**Fig 1F**) and multiple genes that define a homeostatic/resolving macrophage state (*35, 36*) (**Fig S3C**). In stark contrast, UC and CD viromes both robustly promoted inflammatory pathways in macrophages including many cytokines (*IL6*, *IL1B*, *IL8*, *IFNG*) and chemokines (*CCL1, CCL20, CCL24, CXCL3, CXCL5*, **Fig 1G, H**). Gene-set enrichment analysis defined the significant pathways downregulated by non-IBD viromes to be apoptosis, inflammatory and antiviral (**Fig 1I**) while those pathways that were upregulated were pro-survival and pathways known to polarize macrophages toward homeostatic/resolving states such as WNT-β Catenin signaling (**Fig 1I**). In constrast, both UC and CD viromes robustly promoted inflammatory and apoptotic pathways (**Fig 1J,K**). We next confirmed the divergent macrophage activation states induced by non-IBD and IBD viromes using ELISA and qPCR with VLPs isolated from additional patient colon resections. Cytokines IL-10, IL-22, TGFβ or negative regulators of inflammation *CISH* or *SOCS2*, hallmark genes of anti-inflammatory and tissue remodeling type macrophages (*35, 36*) were all consistently and significantly elevated in macrophages exposed to VLPs isolated from 12 different non-IBD patient colon resections (**Fig 1L, Fig S4**). Notably, IL-22 induction is consistent with recent reports of mice mono-associated with apathogenic eukaryotic enteric viruses (*37*). Conversely, macrophages that internalized viromes derived from UC or CD intestinal resections from 8-12 separate patients, repeatedly displayed significantly enhanced production of cytokines TNF, IL-6, Type I and II IFNs, IL-1β and IL-12 that define pro-inflammatory type macrophages as well as other cytokines such as IL-15, IL-8 and IL-31 with designated roles in intestinal inflammation and IBD (*38, 39*) (**Fig 1M, Fig S4**). Importantly, inhibition of endocytosis or heat inactivation of VLPs, failed to induce macrophage cytokine responses (**Fig S5**). However, crosslinking of VLP preparations did not significantly reduce innate responses indicating that virus replication is not an absolute requirement for virome stimulatory capacity (**Fig S5**). Moreover, we observed similar divergent macrophage responses with virus isolated from non-IBD, UC and CD ileum lumen contents (**Fig 1L,M**, **Fig S4, S6, Table S2**). Taken together, these results demonstrate that human enteric viruses residing in colon tissue or ileum content autonomously invoke host innate immune responses. The macrophage response to enteric viruses is atypical to the classical antiviral response normally characterized by the induction of type I IFNs and ISGs. Enteric viruses from a healthy intestine instead dampened the classical antiviral response and promoted anti-inflammatory and pro-survival programs. Conversely, enteric viruses existing within an inflamed UC or CD intestine independently and robustly promoted a state of inflammation.

Intestinal epithelial cells (IECs) express innate immune receptors that recognize and respond to commensal microorganisms. Furthermore, bacteriophages are readily internalized by IECs (*30, 31*). We therefore also tested the ability of healthy or IBD enteric viruses to affect epithelial barrier function and the immune response of IECs. We utilized a trans-well Caco-2 cell monolayer system and exposed IEC monolayers with viruses isolated from non-IBD, UC or CD colon resections and observed significantly more barrier integrity damage induced by both UC and CD-derived enteric viruses in a TLR4-independent manner, as measured by transepithelial electrical resistance (TEER, **Fig S7A,B**). Disrupted barrier integrity elicited by IBD viruses was also associated with a greater downregulation of transcripts for tight junction proteins Zonula occludens-1 (ZO-1) and Occludin (**Fig S7C,D**). Importantly, IFNλ which controls both acute and persistent viral infections in the intestine (*40, 41*) and confers protection from intestinal inflammation (*20*) was preferentially induced by non-IBD viruses (**Fig S7E**). In contrast, viruses isolated from either UC or CD colons promoted production of pro-inflammatory cytokines such as TNF and IL-15 (**Fig S7F,G**) that play crucial roles in intestinal tissue destruction and the pathogenesis of IBD (*38, 39, 42*). Of note, IL-15 was also recently shown to be induced by the murine enteric virome in dendritic cells to promote expansion of innate epithelial lymphocytes (IELs) (*23*). Thus, similar to macrophages, IECs sense and divergently respond to non-IBD and IBD enteric viruses whereby IBD viromes promote both inflammatory responses and disruption of epithelial barrier function.

Fecal microbiota transplants (FMT) to restore disrupted microbiota are proving beneficial in both experimental and clinical settings (*43*) but the virome content of these preparations is currently not considered. We wondered if isolated healthy enteric viruses could overcome the inflammatory response induced by IBD viruses. To this end, we mixed increasing ratios of non-IBD colon resection derived viruses with IBD viruses while maintaining the same overall number of VLPs and exposed human macrophages to them. Increasing amounts of non-IBD viruses were capable of suppressing the ability of IBD-derived viruses to induce pro-inflammatory cytokines such as IL-6, TNF and IL-15 in a dose-dependent manner (**Fig 1N, Fig S8A,B**). Moreover, increasing the ratio of non-IBD viruses could partially restore the ability of macrophages to produce anti-inflammatory cytokines such as IL-22 (**Fig S8C**). To test if the product of the immune response to healthy viruses in the form of secreted soluble factors could also suppress the IBD virome responses, we performed supernatant transfer experiments. In brief, human macrophages were exposed to non-IBD VLPs for 24 hours, then supernatant was collected, clarified, and placed on fresh cells in the presence of IBD colon-derived VLPs. Supernatant from non-IBD VLP treated macrophages was capable of suppressing the production of IL-6, TNF, IL-15 by IBD VLPs (**Fig 1O, Fig S8D, E**). Furthermore, heat inactivation ameliorated suppressive activity of non-IBD virome-induced macrophage supernatant but UV crosslinking did not (**Fig 1O, Supp Fig 8D,E**) demonstrating soluble proteins, but not residual viruses in the supernatant, were eliciting anti-inflammatory activity. Thus, this data demonstrates the potential utility of viruses in a healthy intestine, or the anti-inflammatory immune response to it, to suppress the inflammatory capacity of viruses in a diseased intestine.

To gain a mechanistic understanding of the divergent immunomodulation by non-IBD or IBD derived human colon tissue or ileostomy fluid-derived viromes, we profiled the virus types in each cohort since only fecal VLPs have been examined to date. DNA and RNA were purified from preparations of VLPs from surgical colon resections or ileostomy fluid and characterized by metagenomic sequencing. After filtering out human and other contaminant (primers and adapters) and low-quality data, sequences were assigned a eukaryotic viral or bacteriophage taxonomic lineage. Specific eukaryotic viral families were selected for subsequent analysis based on the alignment length and percent identity for each family that separated the high-quality alignments from potentially spurious alignments (**Table S3**). Total reads aligning to eukaryotic viruses were similar for non-IBD, UC or CD patients, however there was a significant increase in bacteriophage reads in CD patients (**Fig 2A**). Of the high-quality reads for subsequent analysis, we found *Papillomaviridae, Picornoviridae, Coronoviridae, Orthomyxoviridae* and *Anelloviridae* animal viruses, *Virgiviridae* plant viruses and a number of unknown eukaryotic virus families in colon tissue (**Fig 2B**). Furthermore, we found the majority of VLP classifications in human colon tissue to be bacteriophage families consisting of dsDNA *Caudovirales* order including *Siphoviridae, Myoviridae, Podoviridae* and unknown; dsDNA families *Herelleviridae*, *Akkermanniviridae*, *Lipothrixviridae*, as well as ssDNA phage family *Microviridae*. Quantification of viral species richness showed no significant difference between groups (**Fig S9A**) but differential analysis showed a significant elevation in the *Picornoviridae* family member *Enterovirus B,* in both UC and CD colon tissue compared to non-IBD controls (**Fig 2C**). *Enterovirus B* has not been reported in fecal metagenomic studies performed to date, but has been previously detected in resected terminal ileum of immunocompetent pediatric patients with Crohn’s disease (*44*). This further highlights the need to catalog viruses and other microorganisms (*45*) in intestinal tissue that is not fully captured in fecal material. Human enteroviruses are extremely common RNA viruses that spread mainly through the fecal-oral route. Species B enteroviruses consist of coxsackieviruses B1–B6 (CVB1–6), coxsackievirus A9 (CVA9), over 30 serotypes of echoviruses and more than 20 EV-B serotypes that all share the same overall structure found in picornaviruses, a single stranded ∼7500 bp RNA genome. We searched for specific Enterovirus B serotypes in our data and found high identity mapping to *Echovirus B75*, *Echovirus B5* and *Echovirus 26*, that were significantly elevated in both UC and CD colon resection samples. The wide prevalence and persistence of enteroviruses within humans has suggested a potential role for these viruses in complex human disease, including type I diabetes (*46*). Here we now identify *Enterovirus B,* and particularly Echovirus serotypes, as a potential pathogenic factor in UC and CD.

**Fig 2.**
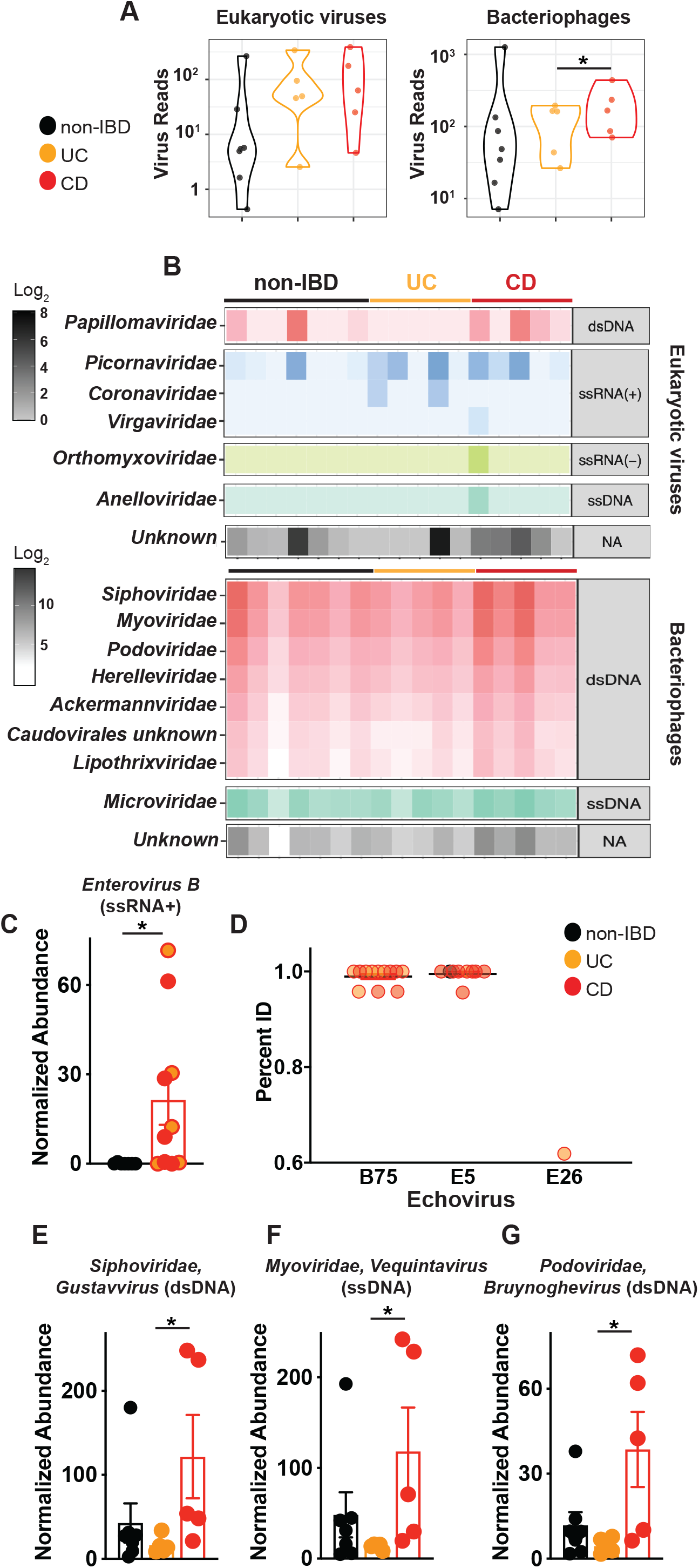
Detection and characterization of VLPs in colon resections from non-IBD, Ulcerative Colitis and Crohn’s disease patients. VLPs were assessed using VLP DNA and RNA metagenomic sequence data. Sequences were assigned a eukaryotic viral or phage taxonomic lineage using primary searches against virus sequence databases and subsequently confirmed in secondary searches using reference databases with additional non-viral taxonomic lineages. Specific eukaryotic viral families were selected for subsequent analysis by gating on the alignment length and percent identity for each family that separated the high-quality alignments from potentially spurious alignments (Table S3). **(A)** Violin plots represent the distribution of individual datasets for total reads aligning to eukaryotic viruses or bacteriophages from non-IBD, UC or CD cohorts; samples were compared using two-sided Wilcoxon signed-rank tests. **(B)** Taxonomic assignments of VLP sequences in non-IBD, UC or CD colon tissue. **(C)** Significantly elevated abundance of eukaryotic virus, *Picornoviridae>Enterovirus B* in UC and CD colon resections. **(D)** Identification of *Echovirus B75*, *Echovirus E5* and *Echovirus E26* Enterovirus B serotypes in UC or CD colon tissue by bbmap. All potential Picornavirales reads were re-mapped one at a time to six Enterovirus B genomes (KF874627.1, NC_001472.1, AF083069.1, AY302539.1, D00627.1, NC_038307.1), using the bbmap tool, mapPacBio.sh with the parameters recommended for remote homologies (vslow k=8 maxindel=200 minratio=0.1). The subset of these reads that mapped to Enterovirus B, along with the previously identified Enterovirus B reads, were plotted, split into Serotypes on the x-axis and percent identity on the y-axis. Each read found is plotted and grouped and colored by the type of sample in which it was found (Non-IBD, UC or CD). Non-IBD versus UC and Non-IBD versus CD were significantly different (p-adj = 0.029, 0.001). **(E)** Top significantly increased genera of *Siphoviridae*, **(F)** *Myoviridae* or **(G)** *Podoviridae* bacteriophages in CD colon resections. Data are the mean of 5-7 biologial replicates. \**P*<0.05, Kruskal–Wallis test with a Dunn’s post-hoc test.

For bacteriophages, CD colon tissue exclusively demonstrated a significant expansion, consistent with reports of CD fecal viromes (*1*). We found significant elevations in total phage aligned reads (**Fig 2A**), a distinct separation of CD from non-IBD and UC by NMDS analysis (**Fig S9B**) and a multitude of significantly increased genus of Caudovirales members *Siphoviridae*, *Myoviridae* and *Podoviridae* (**Fig 2D-F, Fig S9C**). Interestingly, ileostomy fluid VLPs showed significantly reduced viral diversity compared to colon derived VLPs (**Fig S10**) underscoring tissue rather than luminal contents to be the abundant source of virus in the intestine. Detectable eukaryotic families in ileostomy fluid were limited to *Annelloviridae, Papillomaviridae, Picornoviridae* and *Virgiviridae* eukaryotic viruses and the bacteriophage families were *Caudovirales* family members *Siphoviridae, Myoviridae, Podoviridae* and unknown; dsDNA families *Herelleviridae*, *Akkermanniviridae*, *Lipothrixviridae* as well as ssDNA phage family *Microviridae* (**Fig S10**).

We next determined the virus receptor requirement for host responses to human enteric viruses. Virus-derived RNA structures are recognized by pattern recognition receptors (PRRs) such as cytosolic (RIG-I)-like receptors (RLRs), or endosomal Toll-like receptor (TLR)3, 7, 8 and viral DNA is mainly recognized by cyclic GMP-AMP (cGAMP) synthase (cGAS) or TLR9 (*47*). So far, endosomal TLR3, TLR7, TLR9 and cytosolic RIG-I have all been shown to be essential for sensing of the healthy murine virome (*19, 23, 28*). We generated bone marrow–derived macrophages from mice deficient in the RLR signaling adapter mitochondrial antiviral signaling protein (MAVS), DNA sensor cGAS, TLR adapter protein MyD88 or endotoxin sensor TLR4, and exposed them to human colon VLPs (**Fig 3A,B**). Indeed, MAVS, cGAS or MyD88 deletion resulted in attenuated non-IBD virome responses, as measured by pro-survival gene *Cish* (**Fig 3A**). Conversely, deletion of endotoxin receptor TLR4 had little effect (**Fig 3A**), underscoring recognition of virus nucleic acid in VLP-driven responses. Similarly, pro-inflammatory cytokine IL-6 production by colon tissue derived IBD VLPs was also significantly diminished in MAVS-, cGAS-, and MyD88-deficient macrophages but was unchanged in TLR4-deficient macrophages (**Fig. 3B**). Notably, ileostomy fluid-derived VLPs, that had significantly reduced virus diversity compared to colon tissue (**Fig S10**) and was mostly bacteriophages and animal viruses of the *Annelloviridae* family (**Fig S10**), required DNA sensor cGAS exclusively for cytokine induction. Certainly, a dominance of viral-derived DNA was observed in the ileum content whilst the colonic tissue virome was a mix of RNA and DNA (**Fig S11**). This data suggests there is recognition redundancy for normal/non-IBD or IBD enteric viruses but qualitative and quantitative differences in the downstream immune responses induced by them. RLRs RIG-I (retinoic acid-inducible gene I; encoded by *DDX58*) and MDA5 (melanoma differentiation-associated gene 5; encoded by *IFIH1*) are activated by distinct viral RNA structures to differentiate between viral and cellular RNAs. MDA5 senses long double-stranded (ds)RNA, while RIG-I responds to blunt-ended dsRNA bearing a triphosphate (ppp) moiety (*48*). Transfection of purified RNA isolated from non-IBD VLPs, triggered more IL-10 than equal concentrations of RNA from UC or CD VLPs (**Fig 3C**). In contrast, UC or CD VLP RNA triggered more IL-6 production (**Fig 3D**) demonstrating divergent macrophage responses were not due to differences in viral tropism or levels. Importantly, digestion of single-stranded or double-stranded VLP RNA with RNase I or III, respectively, or removal of phosphates with Calf Intestinal Phosphatase (CIP), eliminated this response (**Fig 3C, D**) demonstrating the existence of “non-self” RNA moieties within the human enteric virome and the requirement for both RIG-I and MDA5 in their recognition. Divergent anti- or pro-inflammatory responses by non-IBD or IBD enteric viruses were also induced by transfection of VLP-derived DNA, in a cGAS-dependent manner (**Fig 3E, F**). Thus, altered RNA or DNA moieties that are sensed by host virus receptors RIG-I, MDA-5 or cGAS (and likely other receptors), rather than an adaptive function of viral proteins on the host, contribute to the divergent activation thresholds and immunomodulation by normal/non-IBD and IBD enteric viromes. This deviation in RNA and DNA viral ligands within the diseased virome is attributed to the fluctuations of certain viruses we observed (**Fig 2**). However, it is also appealing to speculate that akin to some bacteria (*49*), normally symbiotic viruses may also become pathobionts when in an inflamed environment.

**Fig 3.**
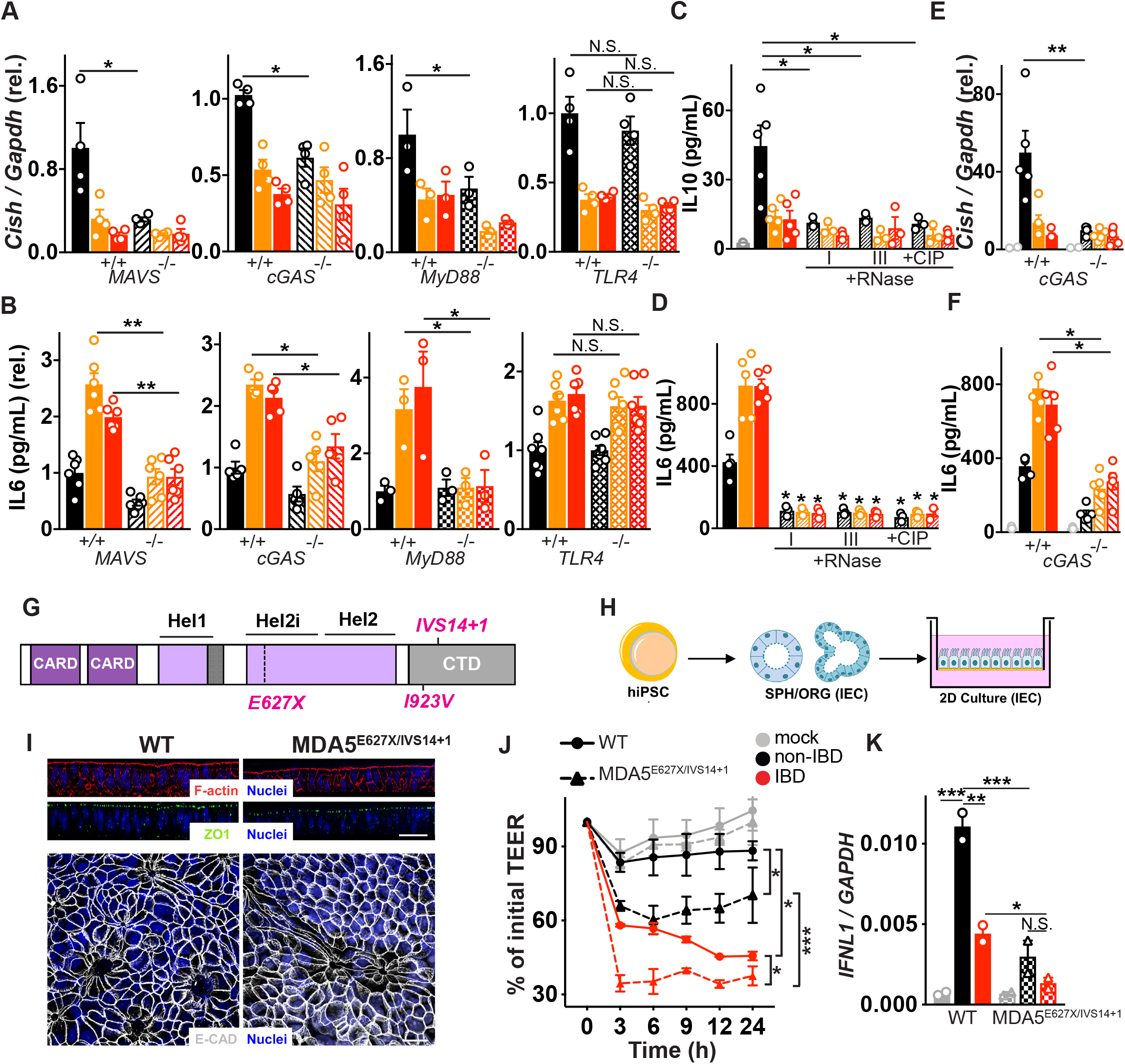
Host viral receptor requirement for sensing human enteric viruses and detrimental consequences for IBD patients with MDA5 loss-of-function variants. (**A**) *Cish* or (**B**) IL-6 levels following exposure of primary bone marrow derived macrophages (BMDMs) from wild-type, MAVS^−/−^, cGAS^−/−^, MyD88^−/−^ or TLR4*^−/−^* mice to non-IBD, Ulcerative Colitis (UC) or Crohn’s disease (CD) colon resection-derived VLPs as measured by qPCR and ELISA. (**C**) IL-10 or (**D**) IL-6 production 24h following transfection of THP1 cells with 2.5 μg of VLP RNA or VLP RNA pre-treated with RNase I (5μg/mL, 1h), RNase III (5μg/mL, 1h) or Calf Intestinal Phosphatase (CIP, 250U, 1h). (**E**) *Cish* levels or (**F**) IL-6 production 24h following transfection of WT or cGAS^−/−^ BMDMs with 2.5 μg of VLP DNA as measured by qPCR or ELISA. (**G**) Schematic of human MDA5 and mapped IBD-associated MDA5 variants rs35744605/E627X, rs35667974/I923V and rs35732034/IVS14+1. **(H)** Schematic of patient lymphoblastoid cell line (LBL) derived human induced pluripotent stem cells (hiPSCs) trans-differentiated into intestinal epithelial cell monolayers. (**I**) Differentiated mature monolayers evaluated for structural integrity markers. Upper panel, orthogonal view of F-actin (Phalloidin, red), tight junction protein ZO1 (green) and nuclei (DAPI, blue) by confocal microscopy. Scale bar represents 50μm. Lower panel, maximal intensity projection images of epithelial adherens junction marker E-cadherin (gray) and nuclei (DAPI, blue). Scale bar is 20 μm. **(J)** Trans-epithelial electrical resistance (TEER) over indicated time or (K) *IFNL1* levels at 24h as measured by qPCR of WT or MDA5 E627Ter/IVS14+1 patient hiPSC-derived intestinal epithelial cell monolayers and exposed to VLPs (10 VLPs/cell) extracted from non-IBD or IBD colon resections. Data are mean ± s.e.m. of 2-5 biological replicates repeated twice. \**P*<0.05, \*\**P*<0.01, \*\*\**P*<0.001, \*\*\*\**P*<0.0001 as determined by one-way ANOVA with Tukey’s multiple comparison test.

Given the existence of genetic mutations within MDA5 that were identified to associate with IBD incidence by GWAS (*29*) and the discovery of elevated *Enterovirus B* of the *Picornoviridae* family in IBD colon tissue, that is exclusively sensed by MDA5 (*50*), we next examined the consequence of these mutations on responses to the human non-IBD and IBD enteric virome. *IFIH1* gain-of-function has been reported as a cause of a type I interferonopathy encompassing a spectrum of autoinflammatory phenotypes including Systemic Lupus Erythematosus and Aicardi-Goutières syndrome (*51*). Yet, intriguingly, all IBD-associated *IFIH1* variants have predicted loss-of-function biological effects, either truncating the protein (rs35744605, E627X), affecting essential splicing positions (rs35732034 in a conserved splice donor site at position +1 in intron 14) or altering a highly conserved amino acid (rs35667974/I923V; **Fig 3G**) within the C-terminal domain (CTD) which recognizes and binds to RNA (*52*). We transfected HEK-293T cells with wild-type MDA5 or E627X or I923V MDA5 coding variants and found the wild-type and MDA5 mutants were expressed at comparable levels although the E627X truncation, that lacks a part of the helicase domain and the entire CTD in which I923 resides, showed slightly reduced MDA5 protein levels (**Fig S12**). Wild-type MDA5 conferred similar responsiveness to the synthetic RNA ligand poly(I:C) and RNA isolated from healthy human colon tissue VLPs but was not responsive to self RNA (**Fig S12**). However, both IBD-associated MDA5 variants exhibited a loss of function phenotype to poly(I:C) (**Fig S12**) in agreement with a previous report (*53*). Importantly, a loss of function phenotype to human colon derived viral RNA was observed for both IBD-associated coding MDA5 variants (**Fig S12**). We next examined the functional consequence of a loss of MDA5 function on intestinal health. MDA5 is predominantly expressed in epithelial cells (**Fig S13**) so we examined functional consequences of loss-of-function MDA5 by converting induced pluripotent stem cells (iPSCs) derived from a compound heterozygote UC patient carrying a single allele of rs35732034/IVS14+1 and a single allele of rs35744605/E627X into human intestinal epithelial cells (hIECs, **Fig 3H**, **Fig S14**). Compared to non-affected individuals, hIECs from patients carrying loss-of-function MDA5^E627X/IVS14+1^ displayed similar morphology and barrier integrity at steady state, as measured by confocal microscopy for F-actin, ZO1 and E-cadherin (**Fig 3I**). However, unlike iPSC-derived hIECs cells bearing wild-type MDA5, MDA5^E627X/IVS14+1^ iPSC-derived hIECs were severely impaired in their ability to maintain barrier integrity when exposed to enteric viruses derived from non-IBD colon resections (**Fig 3J**). Furthermore, damage to MDA5^E627X/IVS14+1^ hIECs was exacerbated when exposed to IBD derived viromes containing elevated *Enterovirus B* (**Fig 3J**). Moreover, IBD-risk MDA5^E627X/IVS14+1^ iPSC-derived hIECs displayed significantly abrogated virome-induced IFNλ production compared to wild-type cells (**Fig 3K**). Thus, abolished innate responses to the virome due to human genetic variation within virus receptor MDA5 has detrimental consequences for intestinal epithelial cell function and integrity. Moreover, intestinal damage is intensified in patients bearing IBD-risk MDA5 variants responding to an IBD virome, underscoring the ability of environmental cues to uniquely impact IBD phenotypes in the context of host genetics.

We next assessed the ability of non-IBD or IBD human colon-resident viruses to shape local immunity and intestinal disease *in vivo*. To achieve this, we first diminished the murine enteric virome of C57BL/6 using an antiviral (AV) cocktail by oral gavage for 10 days (*19*) and then administered human colon resection-derived VLPs 3 times intragastrically (**Fig 4A**). To complement our functional data in macrophages, we first assessed the influence of healthy or diseased enteric viromes on gut resident mononuclear phagocytes (MNPs), especially those bearing the fractalkine receptor CX3CR1 (CX3CR1^+^ MNPs), since they have established roles in the induction of immune responses toward enteric bacteria (*54*) and more recently intestinal fungi (*55*). In parallel, we examined the influence of human enteric viromes on subsets of dendritic cells (DCs) differentially expressing integrins CD11b and CD103 that initiate immune tolerance (*56, 57*). Mice with a healthy humanized virome showed no changes in lamina propria CX3CR1^+^ MNPs compared to wild-type or antiviral treated mice (**Fig 4B,C**) but had elevated tolerizing CD11b^+^CD11c^+^ phagocytes expressing CD103 (**Fig 4B,C, Fig S15**). In contrast, mice with an IBD humanized virome displayed a significant accumulation of MNPs expressing CX3CR1 and a reduction in CD11b^+^CD11c^+^ MNPs expressing CD103 in the colonic lamina propria, both in number and as a fraction of the total CD11b^+^ phagocyte compartment (**Fig 4B,C, Fig S15**). Thus, IBD-associated enteric viruses can promote an inflammatory state in the intestine *in vivo* by spontaneously influencing the ratio of CX3CR1^+^ and CD103^+^ lamina propria MNPs.

**Fig 4.**
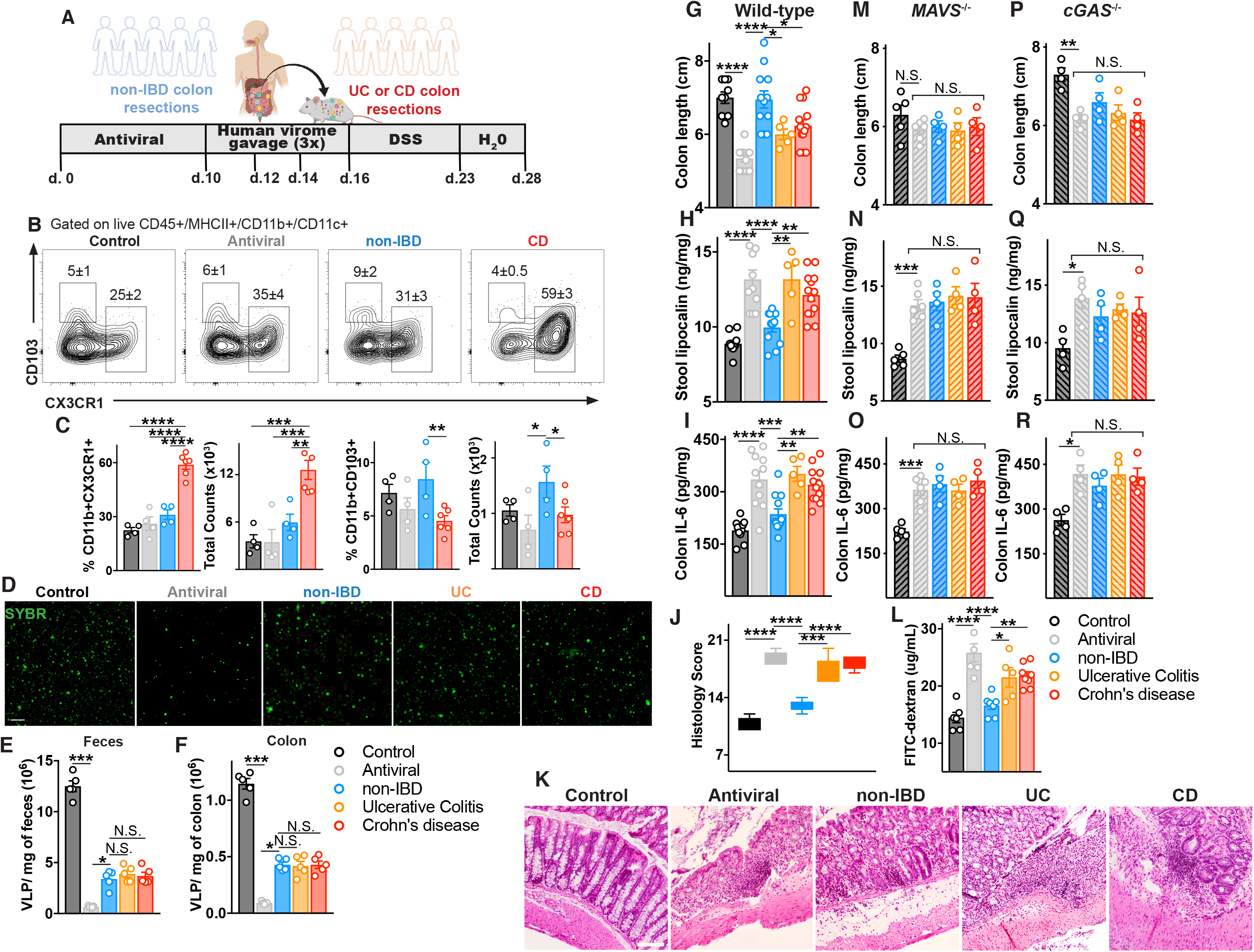
Mice with a “humanized” non-IBD virome had attenuated intestinal inflammation while those with a “humanized” IBD-derived virome exhibited intestinal inflammation. (**A**) Schematic of mouse gut virome depletion using an antiviral cocktail (Acyclovir 20mg/kg, Lamivudine 10mg/kg, Ribavirin 30mg/kg, Oseltamivir 10mg/kg, gray, daily gavage for 10 days), and reconstitution with 200 μL of human non-IBD or IBD VLPs pool (4×10^8^ VLPs per mouse) by gavage on day 10, 12 and 14. 2.5% DSS colitis model was commenced on day 16. (**B**) Flow cytometry plots of CD45^+^MHCII^+^CD11c^+^CD11b^+^ cells from the colonic lamina propria, and percentages and counts of (**C**) CX3CR1^+^ or CD103^+^ mononuclear phagocytes in control, antiviral depleted (AV), non-IBD or IBD humanized virome mice on day 17. (**D-F**) Confocal images and quantification of Viral Like Particles (VLPs) using SYBR Gold in feces or colon tissue of humanized virome mice. (**G**) Colon length, (**H**) stool lipocalin, (**I**) IL-6 levels in colonic explant supernatants cultured for 24 hours following 12 days of induced DSS colitis measured by ELISA. (**J,K**) Representative hematoxylin and eosin–stained sections and blinded histologic scores in mouse colon tissue (scale bar, 50 μm). (**L**) FITC-dextran level in serum 4 hours after oral gavage following 7 days of DSS. **(M-O)** Colon length, stool lipocalin or colonic explant IL-6 levels following induction of DSS colitis in antiviral-treated MAVS−/− mice or **(P-R)** cGAS−/− mice administered human non-IBD, UC or CD colon-derived viromes. Data are mean ± s.e.m. of 4-8 animals. Representative of 2 independent experiments. \**P*<0.05, \*\**P*<0.01, \*\*\**P*<0.001, \*\*\*\**P*<0.0001, two-tailed unpaired t test or one-way ANOVA with Tukey’s multiple comparison test.

The ability of the human enteric virome to contribute to disease phenotype in a dextran sulfate sodium (DSS)-induced colitis model was then assessed. Since this model is more closely associated with UC, mice diminished of their murine virome were administered human enteric viruses derived from colon resections of non-IBD or UC or CD patients 3 times intragastrically (**Fig 4D**). We assessed the abundance of isolated virus-like particles (VLPs) in each treatment group by staining fecal or colon tissue derived VLPs with SYBR. We found a mean of 1.1×10^7^ VLPs/mg of feces and 1.1×10^6^ VLPs/mg of colon tissue in naïve C57BL/6 mice (**Fig 4E,F**). After AV cocktail treatment, fecal VLPs declined ∼20-fold to a mean of 6×10^5^ VLPs/mg in accordance with previous publications in Balb/c mice (*19*) and VLPs in colon tissue declined ∼12 fold to 9×10^4^ VLP/mg. Quantification of viruses in feces or colon tissue of humanized virome mice revealed an equal abundance of ∼4×10^6^ VLPs/mg of feces and ∼4×10^5^ VLPs/mg of colon tissue from mice humanized with healthy, UC or CD viromes (**Fig 4E, F**). Consistent with recent publications (*19, 20*), depletion of viruses in healthy mice lead to significantly exacerbated DSS induced intestinal damage (**Fig 4G-I**). Remarkably, mice repopulated with a healthy humanized virome conferred robust protection against colitis, including less shortening of the colon (**Fig. 4G**), rescued stool lipocalin levels (**Fig. 4H**), rescued colon IL-6 levels (**Fig 4I**). Thus, normal/non-IBD human enteric viromes actively immunodulate and incite protection from intestinal inflammation *in vivo*, consistent with our *in vitro* results. In contrast, both UC or CD human colonic viromes were incapable of rescuing antiviral treated mice and promoted intestinal disease *in vivo* (**Fig 4G-I**), results that were confirmed by histological analysis (**Fig 4J,K**). Moreover, confirming our *in vitro* data, both UC or CD intestinal viromes resulted in reduced intestinal barrier integrity *in vivo*, as measured by gavaged FITC-dextran in the serum (**Fig 4L**). We had demonstrated an autonomous immunodulatory capacity of intestine-derived viromes in cells, however, to assess any virome-microbiome community interactions *vivo*, we performed 16S sequencing of host microbiota. Bacterial diversity and richness were not significantly affected by antiviral treatment or gavage administration of human healthy or IBD viromes (**Fig S16A**). However, in UniFrac-based PCoA analysis, groups were clustered separately dependent on virus depletion or human virome source (P<0.05, PERMANOVA, **Fig S16B**). By proportion (**Fig S16C**) and LEfSe analysis, the bacterial groups most changed by AV cocktail treatment was a reduction in the phyla *Firmicutes* and an elevation certain genera of the *Ruminococcaceae* family (**Fig S16D**). Both healthy and UC humanized virome mice displayed similar changes in murine microbiota (**Fig S16E**) suggesting microbiome changes are unlikely to be the driver of divergent human virome-induced phenotypes. Furthermore, CD humanized virome mice had unique changes in the murine microbiome (**Fig S16F**) reflecting the distinct bacteriophage alterations we had observed in CD colon tissue. Importantly however, virus nucleic acid sensing by the host immune system was an absolute requirement for immunomodulation by healthy or IBD human viromes *in vivo.* Mice deficient in MAVS, the adapter protein of RNA sensors RIG-I and MDA5, or mice deficient in the DNA sensor cGAS, displayed no change in colon length, lipocalin or colon IL-6 production following administration of non-IBD, UC or CD human enteric viromes, compared to antiviral-treated controls (**Fig 4M-R**). Thus, an ability of human enteric viromes to directly shape intestinal disease phenotype *in vivo* was via host innate immune recognition of viruses. Moreover, it demonstrates that fluctuations in the virome are not correlative with disease but rather there are direct functional consequences on host immune and intestinal state due to active host sensing of an altered virome in IBD (*1, 8, 10*) (**Fig 2**)

While disruption of microbiome and virome has been observed in IBD and other diseases, a functional role for the virome has remained elusive. Our data demonstrate that viruses resident to normal human colon tissue are readily endocytosed and elicit an immunomodulatory transcriptional program in macrophages and intestinal epithelial cells, consistently promote production of anti-inflammatory cytokines, and can maintain homeostasis and protect against induced colitis *in vivo*. Conversely, viromes from inflamed intestines, and specifically UC and CD, triggered pro-inflammatory macrophage and IEC barrier loss phenotypes, and intestinal inflammation *in vivo* upon transfer. These functional differences were dependent on innate viral sensors, indicating active and unique recognition of the intestinal virome by the host. Altogether, our findings provide evidence that viruses in a normal/non-inflamed intestine can help build gut immunity in humans - a contrast to the misconception that all viruses are harmful. An alteration in the intestine-resident virome, or disrupted virome sensing due to genetic variation in the virus receptor MDA5, has detrimental consequences for the human intestine. It is appealing to speculate that viruses play a role in the protection mediated by fecal transplants, as well as in the pathology of intestinal inflammation. Therapeutic modulation of the virome through targeted elimination or replacement of disease- or health-driving intestinal viruses presents an novel strategy to treat intestinal and immunological diseases more broadly.

## Acknowledgments

The authors wish to thank the clinical coordinators and patients enrolled in the Prospective Registry in IBD study at Massachusetts General Hospital (PRISM). We would also like to thank the MGH Next Gen Sequencing core, Thomas Diefenbach at the Ragon Institute Imaging Core for confocal imaging of VLPs, the Harvard Medical School electron microscopy facility, Hembly Rivas (Harvard Virology Ph.D. program) for assistance with MDA5 experiments, Laurie Pruneau and Marie-Eve Rivard (Montreal Heart Institute) for technical expertise with iPSC to human epithelial cell trans-differentiation using LCL cells lines from NIDDK Inflammatory Bowel Disease Genetics Consortium (IBDGC) and Chloé Lévesque for confocal immunofluorescence analyses of the hiPSC-derived epithelial 2D cultures. The NIDDK IBDGC was conducted by the IBDGC Investigators and supported by the National Institute of Diabetes and Digestive and Kidney Diseases (NIDDK). The lymphoblastoid cell lines and related data from the IBDGC reported here were supplied by the NIDDK Central Repositories. Other than J.D.R., this manuscript was not prepared in collaboration with investigators of the IBDGC study and does not necessarily reflect the opinions or views of the IBDGC study, the NIDDK Central Repositories, or the NIDDK. We thank Amy Avery (Massachusetts General Hospital) for technical assistance with 16S sequencing and Raza Hoda (Massachusetts General Hospital) for blindly scoring intestinal pathology. We thank Corinne Maurice (McGill University), Irah King (McGill University), Naama Geeva-Zatorsky (Technion) and the entire Jeffrey lab for valuable scientific discussions and Robert Anthony, James Moon and Ramnik Xavier for critical reading of the manuscript.

## Funding

This study was supported by the Kenneth Rainin Foundation (Innovator and Synergy Awards to K.L.J), NIH R21AI144877 (K.L.J), NIH R01DK119996 (K.L.J), Harvard Catalyst | The Harvard Clinical and Translational Science Center (National Center for Advancing Translational Sciences, National Institutes of Health Award UL 1TR002541) and financial contributions from Harvard University and its affiliated academic healthcare centers (K.L.J), Canadian Institute of Health Research (CIHR) postdoctoral fellowship (H.A.), NIH P30DK040561 (R.S.), Canada Research Chair (J.D.R), NIH DK062432 (J.D.R) and NIH RC2DK116713 (S.H.). K.L.J is a John Lawrence MGH Research Scholar 2020-2025.

## Author contributions

F.A. conducted and analyzed the majority of *in vitro* and *in vivo* experiments, functional assays, developed methods, and generated resources; H.A. assisted with and analyzed VLP uptake flow cytometry and assisted with DSS models, S.D. performed RNA sequencing experiments and performed and analyzed lamina propria flow cytometry experiments with help from H.A.; T.S. assisted with *in vivo* and *in vitro* assays, L-P.W and R.S. analyzed RNA sequencing data; R-U.R. analyzed 16S sequencing; J.P., P.G and J.D.R. provided lymphoblastoid cell lines (LCL) from MDA5 variant IBD patients and performed LCL to iPSC to intestinal epithelial trans-differentiation of these cells., R.H. provided clinical intestinal and ileostomy samples post-surgery; K.M and S.H. analyzed VLP sequencing; K.L.J conceived and supervised the study, acquired funding and wrote the final manuscript.

## Competing interests

The authors declare no competing interests.

## Additional information

### Author Information

Correspondence and requests for materials should be addressed to Kate L. Jeffrey

### Data and materials availability

The data presented in this study are tabulated in the main text and <u>supplementary materials</u>. RNA sequencing raw and processed data has been deposited at Gene Expression Omnibus (GEO) and is accessible using accession number GSE135223. Human colon resection and ileostomy fluid virus like particle (VLP) sequencing are deposited at European Nucleotide Archive (ENA) under accession number PRJEB44121 and 16S rRNA of murine feces of humanized virome mice are deposited at Sequencing Read Archive (SRA) under accession numbers SUB9443038.

## Supplementary Materials for

### Materials and Methods

#### Patient intestinal resections and ileostomy fluid

All human samples were collected under Institutional Review Board (IRB)-approved protocols by Massachusetts General Hospital (MGH) and Research Blood Components, including informed consent obtained in accordance with relevant ethical regulations. Fresh colon resections were collected from patients that were enrolled in the “Prospective Registry in IBD Study at Massachusetts General Hospital (PRISM, IRB# FWA00003136, **Table S1**). Study research coordinators obtained consent, and medical history was obtained and confirmed by review of the electronic medical record. Inflammation in UC or CD colon resection samples was confirmed grossly and microscopically by a clinical pathologist. Non-IBD controls included individuals undergoing colon resection surgeries for colorectal cancer, familial adenomatous polyposis (FAP) or Diverticulitis. Lack of inflammation in non-IBD control resection samples was confirmed by a clinical pathologist. Ileostomy fluid was collected from patients enrolled in the “Quantifying the Pro-Inflammatory Nature of Human Fluid Collection Study” that was approved by the Committee for the Protection of Human Subjects at MGH (IRB#2011P002755, **Table S2**).

#### Virus Like Particle (VLP) Isolation

100 mg of colon tissue or centrifuged ileostomy fluid was resuspended in 500 μL of saline magnesium buffer (SM) buffer (8 mM MgSO_4_, 100 mM NaCl, 50 mM Tris-HCl pH 7.4, and 0.002% (w/v) gelatin, passed through a 0.02μm Whatman filter), homogenized using sterile metal beads, and centrifuged. The supernatant was then passed through a 0.45 μm pore-sized membrane, followed by 0.22 μm pore-sized membrane (Millipore). The filtrates were treated with lysozyme (10 μg/mL; Sigma) for 30 min at 37 °C, followed by incubation with 0.2 volumes of chloroform for 10 min to degrade any remaining bacterial and host cell membranes and then centrifuged at 2,500 × g for 5 min at room temperature. The aqueous phase was collected and incubated with DNase I (3U/200μL, Sigma) for 1 h at 37 °C to remove any non-virus protected DNA. Enzyme activity was inactivated by incubation at 65°C for 15 min as described (*1*). To remove any potential residual endotoxin, samples were treated with Polymyxin B (10 μg/mL, Sigma) for 30 minutes. Undetectable endotoxin was confirmed using Limulus Amebocyte Lysate (LAL) assay kit (GenScript) following the manufacturer’s instructions.

#### Quantification of VLPs

Virus-like particles (VLPs) were diluted 10-fold serially, stained for 30 minutes with 10× SYBR Gold (Thermo Fisher Scientific) for nucleic acid or Dialkylcarbocyanine (DiI, Thermo Fisher Scientific) for lipid bilayers of enveloped viruses, and imaged using a Zeiss LSM510 laser scanning confocal microscope. Images were captured using Zeiss software (ZEN). Particles <0.5 μm in diameter were regarded as VLPs. SM buffer was used as a negative control. 10 VLP images were captured per sample and VLPs were counted using an image analyzer (InnerView™).

#### VLP sequencing

Total RNA and DNA was extracted from VLPs on a MagNA Pure 24 instrument (Roche), according to the manufacturer’s instructions. In order to evaluate samples for both RNA and DNA viruses, the total nucleic acids were randomly amplified as described previously (*2, 3*) using barcoded primers consisting of a base-balanced 16-nucleotide-specific sequence upstream of a random 15-mer and used for NEBNext library construction (New England BioLabs). The libraries were multiplexed on an Illumina NextSeq (Washington University Center for Genome Sciences) using the paired-end 2 × 150 protocol.

#### Identification and analysis of viral-like sequences

Unprocessed paired-end reads were processed through a multistage quality-control procedure to remove primers and adapters, human and other contaminant and low-quality sequence data. Exact duplicate sequences were removed reserving a single copy and all remaining sequence dereplicated allowing for 4 substitutions. These high-quality / low-redundancy sequences were systematically queried against protein or genomic reference databases using MMseqs2 (*4*) translated and untranslated search strategies. Sequences assigned a eukaryotic viral or phage taxonomic lineage were first identified using primary searches against virus sequence databases and subsequently confirmed in secondary searches using reference databases with additional non-viral taxonomic lineages. Specific eukaryotic viral families were selected for subsequent analysis by gating on the alignment length and percent identity for each family that separated the high-quality alignments from potentially spurious alignments (**Table S3**). Only high-quality reads were used for subsequent analysis. Specific phage reads were selected for subsequent analysis by gating on an alignment length of 75 nt, to account for the current state of phage taxonomy. In order to determine which phage and eukaryotic viral reads are differentially abundant in the three cohorts, we analyzed the reads at the family, genus and species levels. The non-parametric Kruskal–Wallis test was used to assess the difference in median abundance of eukaryotic viral and phage taxa in the Non-IBD, UC and CD cohorts. The Dunn’s post-hoc test was used to correct for multiple comparisons and only those taxa with an adjusted p-value < 0.05 were considered significant. The mean abundance of viral or phage reads, scaled by library size, was determined for each sample in the colon and ileostomy samples. The non-parametric Welch’s paired t-test was used to determine which cohorts had significantly different numbers of viral or phage reads.

#### Human peripheral blood derived macrophages

Peripheral Blood Mononuclear Cells (PBMCs) were isolated from 20-30 mL blood buffy coats from healthy human volunteers (MGH blood components lab). Briefly, mononuclear cells were isolated by density gradient centrifugation of PBS-diluted buffy coat/blood (1:2) over Ficoll-Paque Plus (GE Healthcare). The PBMC layer was carefully removed and washed 3 times with PBS, mononuclear cells were collected as PBMC. In order to obtain macrophages, PBMCs were re-suspended in X-VIVO medium (Lonza) containing 1% penicillin/streptomycin (Gibco) and incubated at 37°C, 5% CO_2_ for 1h to adhere to the tissue culture dish. After 1h, adherent cells were washed 3 times with PBS and differentiated in complete X-VIVO medium containing 100 ng/mL human M-CSF (Peprotech) for 7 days at 37°C, 5% CO_2_. On day 4, cultures were supplemented with one volume of complete X-VIVO medium containing 100 ng/mL human M-CSF.

#### Delivery of VLPs to macrophages

VLPs or SM buffer (8 mM MgSO_4_, 100 mM NaCl, 50 mM Tris-HCl, pH 7.4, and 0.002%, w/v, gelatin) were pre-treated for 30 minutes with Polymyxin B (Sigma) to ensure endotoxin-free conditions. Media was removed from 1×10^6^ adherent macrophages and VLPs (10 VLPs per cell) or SM buffer (“mock”) was added drop-wise onto the cells at a fixed volume sufficient to cover the well surface. VLPs were allowed to adsorb to adherent cells for 1 hour at 37°C with occasional rocking. After 1 hour, VLPs were aspirated, cells were gently washed with PBS, and media was replaced (this was considered time = 0). For heat inactivation experiments, VLPs were first inactivated for 1 hour at 60°C. For UV crosslinking of VLPs, VLPs were placed in a UV Stratalinker 2400 from Stratagene on ice to UVB-irradiate three times with 200 mJ/cm2 for 15 minutes. For *in vitro* ratio experiments, non-IBD colon resection-derived VLPs were mixed with CD colon resection-derived VLPs at increasing ratios to the final concentration of 10 VLPs per cell and VLPs were delivered to macrophages as described above. For supernatant *in vitro* experiments, the supernatant from macrophages that were exposed to the non-IBD VLP 24 hours prior was mixed with IBD VLPs and added drop-wise onto the cells. VLPs and supernatant mix were allowed to adsorb to adherent cells for 1 hour at 37°C with occasional rocking. Heat inactivated (95°C for 10 min) or UV crosslinked (three times 200 mJ/cm2 for 15 minutes) supernatant were used as controls.

#### Fluorescent labeling of VLPs and uptake assays

Purified, quantified colon resection derived VLP preparations were labeled with 0.01 μM SYTO^TM^ 16 Green Fluorescent Nucleic Acid Stain (Thermo Fisher Scientific, #S7578) according to manufacturer’s instructions. Labeled VLPs were then separated from unincorporated dye using PD10 Sephadex G-25 desalting columns (GE Healthcare, #17085101) according to manufacturer’s instructions. Uptake assays of SYTO16-labeled VLPs were performed with human peripheral blood monocyte derived macrophages. Cells were seeded at a density of 10^5^ cells/well in a 24-well plate. Cells were delivered 10^6^ or 10^7^ non-IBD, UC or CD VLPs or SM buffer control by adsorption in 100 μL for 1h, washed and left for 3 hours at 37°C for uptake analysis or 4°C for adsorption analysis. Cells were removed from the culture plate surface with cold PBS, washed in flow cytometry buffer twice, stained for live–dead discrimination and acquired on an LSR II flow cytometer (BD Biosciences). For the inhibition assays, cells were treated with 3.0 mg/mL of Brefeldin A (Invitrogen, #00-4506) for 30 min at 37°C before addition of labeled VLPs and during the adsorption. Data was analyzed using FlowJo v10 software (TreeStar).

#### RNA sequencing and analysis

Poly(A)-mRNA was enriched using the NEBNext Poly(A) mRNA magnetic isolation module (NEB), according to manufacturer’s instructions. Libraries were then generated using the NEBNext Ultra II RNA Library Prep Kit for Illumina (NEB). To allow for multiplexing of samples, libraries were dual indexed using indices from NEBnext Multiplex Oligos for Illumina (NEB). Paired-end sequencing was performed using a NovaSeq 6000. Samples were run in replicate across three separate lanes. Transcriptome mapping was performed with STAR version 2.5.4a(*5*) using the NCBI37 assembly gene annotations. Read counts for individual genes were produced using the unstranded count feature in HTSeq 0.9.1(*6*). Differential expression analysis was performed using the EdgeR package(*7*) after normalizing read counts and including only those genes with count per million reads (cpm) greater than 1 for one or more samples(*8*). Differentially expressed genes (DEG) were defined based on the criteria of minimum 2-fold change in expression value and false discovery rate (FDR) less than 0.05. Gene Set Enrichment Analysis (GSEA) (*9*) was adopted for functional annotation analysis.

#### VLP RNA or DNA isolation

For VLP RNA or DNA transfection experiments, VLP preparations were mixed with Trizol-LS (Invitrogen) and RNA and DNA fractions isolated. DNA was further purified by ethanol precipitation. The RNA fraction was subjected to RNeasy Mini Kit (Qiagen) with on-column DNase I digest for clean-up and removal of residual DNA. For enzyme treatments of nucleic acids, 2.5 μg of RNA was treated with RNase I (5 μg/mL, Ambion) or RNase III (5 μg/mL, Ambion) or with Calf Intestinal Phosphatase (CIP, 250U, Roche) at 37 °C for 1 h. Enzyme-treated RNA was purified with RNeasy Mini Kit (Qiagen) before transfection. Human macrophages were transfected with 2.5 μg of RNA or DNA using Lipofectamine 2000 (Thermo Fisher Scientific) according to the manufacturer’s protocol.

#### ELISA

Cell culture supernatants were removed from human macrophages and centrifuged to remove debris and non-adherent cells. Secreted cytokines were measured using a human Cytokine 25-Plex Magnetic Panel for Luminex® Platform (ThermoFisher, EPX250-12166-901) and read on a Luminex FLEXMAP 3D® instrument. Human and mouse TNF, IL-6, IFN-*β*, IL-8, IL-15, IL-10 and IL-22 ELISA kits (R&D) were also used. Supernatants and kit-supplied cytokine standards were processed in triplicate according to the manufacturer’s instruction.

#### Quantitative PCR

RNA was extracted using the RNeasy Mini Kit (Qiagen) with on-column DNase digest (Qiagen) according to manufacturer’s instruction. 100ng-1µg RNA was used to synthesize cDNA by reverse transcription using the iScript cDNA Synthesis Kit (Bio-Rad). Quantitative PCR reactions were run with cDNA template in the presence of 0.625µM forward and reverse primer and 1x solution of iTaq Universal SYBR Green Supermix (Bio-Rad). Quantification of transcript was normalized to the indicated housekeeping gene. A complete list of primer sequences is provided in Supplementary Table 4.

#### Human Caco2 cells monolayer trans-well model

Human Caco2 epithelial cells were purchased from the American Type Culture Collection (ATCC) and maintained in DMEM with 10% heat-inactivated fetal bovine serum (FBS) and 1% Penicillin-Streptomycin (P/S) solution. Toll-like Receptor 4 (TLR4)-null Caco2 cells were kindly provided by Dr. Richard Hodin’s lab (Massachusetts General Hospital). For creating the monolayer, wild-type or TLR4-null Caco2 cells were seeded onto collagen-coated polyethylene terephthalate (PET) filter supports (1 μm pore size, Corning) at a concentration of 3×10^5^ cells/well and were incubated (37°C, 5% CO_2_) for 2-3 weeks. Both the apical and the basolateral side of the trans-wells were fed with DMEM supplemented with 10% FBS and 1% P/S solution. After 2-3 weeks the integrity of the monolayer was measured by transepithelial electrical resistance measurements (TEER) using the Millicell-ERS electrical resistance measuring system (Millipore). The electrodes were immersed in a way that the shorter electrode was in the inner well and the longer electrode was in the outer well. 200-220 Ω.cm^2^ resistance was indicated as a confluent monolayer. When TEER peak was reached, and maintained for 3 days, the monolayer was used for subsequent experiments. SM buffer and LPS (100 ng/mL) were used as negative and positive controls, respectively. Apical compartment surface was incubated with VLPs whereby VLPs were allowed to adsorb to adherent cells for 1 hour at 37°C (10 VLPs per cell). After 1 hour, VLPs were aspirated, cells were gently washed with PBS, and fresh media was replaced (this was considered time = 0). The cells were then returned to the cell culture incubator (37°C, 5% CO_2_) and TEER measured every 6 hours up to 48 hours. All experiments were performed in triplicate.

#### HEK293T IFNβ luciferase reporter assay

HEK293T cells were maintained in 48-well plates in Dulbecco’s modified Eagle medium (Cellgro) supplemented with 10% FBS and 1% penicillin/streptomycin. Cells were transfected with 10 ng of indicated FLAG-MDA5 plasmids (kindly provided by the T. Fujita, Kyoto University), *IFNB*-promoter driven firefly luciferase reporter plasmid (100 ng) and a constitutively expressed Renilla luciferase reporter plasmid (pRL-CMV, 10 ng) by using Lipofectamine2000 (Life Technologies), according to the manufacturer’s protocol. Empty vector was used to maintain a constant total amount of DNA and lipofectamine. Media was changed 6 hours after the first transfection and cells were additionally transfected with 1μg/mL of RNA isolated from either non-IBD virus like particles (VLPs), polyinosinic:polycytidylic acid (poly I:C, Sigma), or RNA isolated from the cytoplasm of HEK-293T cells (self RNA). Cells were lysed 24 h post-stimulation and *IFNβ* promoter activity was measured using the Dual Luciferase Reporter assay (Promega). Firefly luciferase activity was normalized against Renilla luciferase activity.

#### Differentiation of human iPSC into intestinal epithelial cells

Two lymphoblastoid cell lines (LCL) were selected from the NIDDK IBDGC Repository based on their *IFIH1* genotypes: (1) an LCL line carrying wild-type *IFIH1* alleles from a 39 year-old Female Caucasian and (2) an LCL line that is a compound heterozygote for rs35732034 (IVS14+1) and rs35744605 (E627X) from a Caucasian 47 year-old male Ulcerative Colitis patient. Specifically, the risk allele at rs35732034 (IVS14+1) is a splice donor site substitution at position +1 in intron 14 that leads to exon 14 skipping and the truncation of the C-terminal domain, and the risk allele at rs35744605 (E627X) is a nonsense variant leading to the loss of the C-terminal half of the protein (*10, 11*). The two LCL lines were then reprogrammed to human induced pluripotent stem cells (iPSCs) as described (*12*) via nucleofection with four episomal reprogramming plasmids (pCE-hUL, pCE-hSK, pCE-hOCT3/4, and pCE-mp53DD) and selection of TRA-1-81 positive reprogrammed hiPSC colonies. LCL-derived hiPSC were then used to generate polarized epithelial monolayers using a stepwise differentiation protocol involving a conversion to definitive endoderm (DE) using the STEMdiff Definitive Endoderm kit (StemCell Technologies). This was followed by the formation of hindgut (HG) structures with a treatment of 3 μM CHIR99021 and 250 ng/ml FGF4 for 3-4 days, as previously described (*13*). HG structures were harvested and then centrifuged at 100g for 5 min, embedded in Matrigel (Corning) and cultured for 15 days in Intestinal Growth Medium with feeding every 2-3 days as described (*14*) but using human recombinant inducers rhWnt3a (Cedarlane), rhEGF (Cedarlane), rhNoggin (Cedarlane), rhR-Spondin-1 (Millipore-Sigma) and rhHGF (Cedarlane). Following this, the spheroids were cultivated in human organoid culture medium (HOCM) and passaged every 6-8 days for maturation into 3D Intestinal epithelial cell (IEC) cultures of spheroids (SPH) and organoids (ORG) as described (*13, 14*). Intestinal epithelial cell structure and function was then assessed (see Supplementary Fig 14).

#### Delivery of Virus like Particles (VLPs) to hiPSC-derived intestinal epithelial cell monolayers

Spheroids were recovered from the Matrigel by treatment with Cell Recovery Solution (Corning), dissociated in TrypLE Express for 20 min at 37C and further dissociated by pipetting. A typical confluent Matrigel drop of spheroid was used for five Transwell inserts (about 2 × 10^5^ cells). The cell suspensions were centrifuged at 450g for 5 min and resuspended in HOCM (100 μL per Transwell). Cells were seeded on the apical side (upper chamber) of the collagen-coated Transwell unit (PTFE COL 0.4 μm 0.33 cm^2^, Corning) and 600 μL of HOCM was added to the bottom chamber. Transepithelial Electrical Resistance was measured every 2-3 days using Epithelial Voltohmmeter (EVOM2, World Precision Instruments, LLC) until a plateau was reached (similar TEER value on three consecutive measurements). Cells on the apical compartment surface were infected with VLPs (MOI 10) as described above. VLPs were allowed to adsorb to adherent cells for 1 hour at 37°C. After 1 hour, VLPs were aspirated, cells were washed with PBS, and fresh media was replaced. SM buffer and poly I:C (50 μg/mL) were used as negative and positive controls, respectively. The cells were then returned to the cell culture incubator (37°C, 5% CO_2_) and the integrity of the monolayer was measured by transepithelial electrical resistance measurements (TEER) every 6 hours up to 24 hours.

#### MDA5 expression in epithelial cells

##### Western Blot

Mouse intestines were harvested and pre-digested in 1µM ethylenediaminetetraacetic acid (EDTA) and 3µM dithiothreitol (DTT) at 37°C for 15 minutes with shaking (400 rpm) to isolate epithelial cells. Remaining tissue was minced and digested with collagenase type II (Gibco) for 45 minutes at 37°C with constant agitation followed by Percoll density-gradient centrifugation to isolate single cells from the lamina propria. Cells were incubated with biotinylated anti-CD45 to purify the hematopoietic cell population. Beads were gently washed, and bound cells isolated by MACS mini-column purification using streptavidin magnetic beads. Epithelial cells and lamina propria hematopoietic cells were lysed in 200 μL of RIPA buffer (Tris-HCl, pH 7.4, 50 mM, NaCl 150mM, NP-40 1%, sodium deoxycholate 0.5%, SDS 0.1%) with a cocktail of protease and phosphatase inhibitors (1%, Roche). 20 μg of protein was loaded per lane and separated on a 4-12% SDS/PAGE gel and transferred to PVDF. Ponceau stain (Sigma) confirmed protein transfer, and the membrane was blocked for 1 h at 25°C with 5% skim milk in TBS/0.1% Tween-20 (TBS-T), and then incubated overnight at 4°C with the indicated primary antibodies (diluted 1:1000 v/v in TBS-T with 3% BSA). Primary antibody was washed off and the membrane was incubated with the indicated secondary antibodies (diluted 1:10000 in TBS-T). The following primary antibodies were used: anti-MDA5 antibody (Sigma), *β*-actin (Sigma) was used as a loading control, and Villin (Santa Cruz) was used to confirm purity of epithelial cells.

##### Immunofluorescence microscopy

The ileum was fixed with 4% paraformaldehyde. After antigen retrieval (Sigma, Citrate Buffer, #C9999), sections were treated with autofluorescence blocking MaxBlock, blocked in 5% normal goat serum with 1% BSA in PBS (Blocking buffer) for 1 hour at RT, and stained with the primary antibody anti-MDA5 (Sigma; #SAB3500356) overnight at 4°C at a final concentration of 1:500. Slices were washed and incubated with goat anti-rabbit secondary antibody for 1 hour at RT at a final concentration of 1:500. The slices were washed, counterstained with DAPI (Vector), and mounted for confocal microscopy analysis. All images were collected using a confocal microscope (Nikon A1) (400×).

#### Mice

Five- to six-week-old female C57BL/6 mice, MAVS−/−, cGAS−/− or TLR4−/− (C3H/HeJ) were purchased from The Jackson Laboratory. All mice were housed in specific pathogen-free conditions according to the National Institutes of Health (NIH), and all animal experiments were conducted under protocols approved by the MGH Institutional Animal Care and Use Committee (IACUC), and in compliance with appropriate ethical regulations. For all experiments, age-matched mice were randomized and allocated to experimental group, with 5-8 mice per group, and repeated three independent times. No statistical method was used to determine sample size. For “humanized” virome mice, mice were first depleted of viruses using an antiviral cocktail (Acyclovir 20mg/kg, Lamivudine 10mg/kg, Ribavirin 30mg/kg, Oseltamivir 10mg/kg) by daily gavage for 10 days as described (*15*). AV-treated mice were then gavaged with 200 μL of SM buffer as a control or 4×10^8^ VLPs isolated from non-IBD, UC or CD patient fresh colon resections. Mice received the human VLP gavage every other day for three times in total. To ensure that the gastric pH would not affect the viability of the gavaged VLPs, each mouse was gavaged with 100 μL of 1M NaHCO_3_ 15 min prior to VLP gavage to keep gastric pH at 7.4. Quantification of viruses in different treatment groups was performed by confocal imaging of VLPs isolated from feces and colon tissue.

#### Bone marrow-derived macrophages

Bone marrow-derived macrophages (BMDMs) were produced from MAVS-deficient, cGAS-deficient or TLR4-deficient mice (obtained from The Jackson Laboratory) and their respective littermate controls. Briefly, bone marrow was flushed from tibia and femur and allowed to adhere to a non-treated tissue culture plate for 1 day. Non-adherent cells were then differentiated to macrophages in DMEM containing 10% FBS, 1% L-Glutamine, 1% penicillin/streptomycin, 0.1% *β*-mercaptoethanol, 5 ng/mL of interleukin 3 (IL-3, Peprotech) and Macrophage Colony Stimulating Factor (M-CSF, Peprotech) for 7 days. Macrophage maturity was assessed by surface expression of CD11b and F4/80 with flow cytometry. MyD88−/− immortalized macrophages were obtained from Kate Fitzgerald (UMass Medical School).

#### Isolation of lamina propria cells for flow cytometry

Colons were harvested from mice and caeca were removed. The tissue was placed in RPMI containing 5% fetal bovine serum (FBS) and fat was removed by careful dissection followed by gentle rolling along a moist paper towel. The tissue was washed gently then inverted and rinsed to remove loose fat and fecal matter. Pieces of the colon were transferred to a solution of RPMI containing 2% FBS, 1µM ethylenediaminetetraacetic acid (EDTA), 3µM dithiothreitol (DTT) and incubated at 37°C for 15 minutes with shaking (400 rpm). Next, tissue pieces were rinsed with fresh FBS-containing media to wash off EDTA/DTT and tissue was again rolled gently along a moist paper towel. Tissue was cut into very small pieces and transferred to a digestion medium: each sample was placed in 25mL of RPMI media containing 2% FBS and mixed with 12.5mg Dispase II (Gibco) and 37.5mg type IV collagenase (Worthington). Digestion was allowed to take place for 45 minutes at 37°C with constant agitation, and halfway through this period, tissue was disrupted further by pipetting. After 45 minutes, the digestion solution was pipetted again and smashed through a 100µm mesh. Digestion was quenched by addition of 25 mL of RPMI containing 5% FBS. The suspension was spun down for 10 minutes at 450g and the resulting pellet was resuspended in fresh media, passed through a 40 µm mesh, washed with additional fresh media and spun down again. The final pellet represented the lamina propria fraction of the colon. Cells were blocked, stained, and washed for acquisition on an LSRII flow cytometer (BD). Antibodies used for flow cytometry can be found in Supplementary Table 5. All data was analyzed using FlowJo v10 software (TreeStar).

#### Dextran Sodium Sulfate induced colitis (DSS)

Mice were administered 2.5% Dextran Sodium Sulfate Salt (DSS, MW = 36,000-50,000 Da; MP Biomedicals) in drinking water ad libitum for 7 days (freshly prepared every other day), followed by regular drinking water for 5 days. Mice were sacrificed at day 12. Colon length was measured and fecal samples were homogenized in PBS, spun down and supernatants were collected and assayed for Lipocalin-2 by ELISA according to manufacturer’s instructions (R&D). To assess intestinal permeability, mice were given FITC-dextran tracer (Sigma, 4 kDa, 0.6 mg/g body weight) intragastrically in 0.2 mL PBS and FITC levels were measured 4h later in hemolysis-free serum using a Fluorescence Spectrophotometer. For cytokine measurements, day 12 colon tissue (approximately 1 cm) was washed in PBS and then placed in a 24 well plate containing 750 μL of RPMI medium with 1% penicillin/streptomycin and incubated at 37°C with 5% CO_2_ for 24 hours. Supernatants were collected and centrifuged for 10 min at 4°C and IL-6 levels were assessed by ELISA according to manufacturer’s instructions (R&D). For histology, tissue was collected from the distal part of the colon and hematoxylin and eosin staining was performed. Histologic evaluation performed in a blinded fashion and scored based on the criteria outlined previously (*16*). Three main categories were used to reflect the severity of histopathology: (i) inflammatory cell infiltrates, (ii) epithelial changes and (iii) mucosal architecture.

#### 16S sequencing and analysis

QIAamp DNA Stool Mini Kit was used to extract DNA from feces. The hypervariable region (V4) of the 16S rRNA gene sequences were amplified by using PCR with adaptor primer set containing 12bp multiplex identifier sequences (Integrated DNA Technologies, Coralville, IA). Each reaction contained template DNA, each primer (0.2 μM), and 2× Platinum™ Hot Start PCR Master Mix (Invitrogen, Carlsbad, CA). PCR conditions were as follows: 94°C for 3 min, followed by 22 cycles at 94°C for 45 sec, 52°C for 1 min, and 72°C for 1.5 min and a final extension step at 72°C for 10 min. Triplicate reactions were prepared, pooled, and purified using SpeedBead™ carboxylate magnetic bead solution (GE Healthcare, Marlborough, MA). The amplicons were quantified using KAPA Library Quantitation kit (KAPA, Cape Town, South Africa) and an equimolar amount of each sample was sequenced on the MiSeq System (Illumina, San Diego, CA) with 2×150 paired-end run parameters. Microbiome bioinformatics analysis was performed with QIIME 2 release 2018.11 (*17*). In brief, cutadapt was used to trim 515F/806R primers in the paired end sequences. Subsequently, the sequences were stripped of noise by using the DADA2 (*18*) denoiser program. The denoised, trimmed and higher quality amplicon sequence variants (ASVs) were aligned with MAFFT (*19*) and used to construct a phylogeny with fasttree2 (*20*). Alpha-diversity metrics (Shannon’s diversity index), beta diversity metrics (UniFrac (*21*)) and Principle Coordinate Analysis (PCoA) were estimated using q2-diversity after samples were rarefied (subsampled without replacement) to 69009 sequences per sample. All ASVs were assigned taxonomy using the q2-feature-classifier (*22*). In brief, a naïve Bayes classifier was trained on the Greengenes 13_8 99% OTUs reference sequences (*23*) to assign taxonomy to each ASV. The compositional differences among the groups were determined by a linear discriminant analysis using LEfSe (*24*) with a threshold of 2.0 on the logarithmic score using Galaxy (https://huttenhower.sph.harvard.edu/galaxy/).

#### Statistical analysis

Results are shown as mean ± s.e.m. Visual examination of the data distribution as well as normality testing demonstrated that all variables appeared to be normally distributed. Comparisons and statistical tests were performed as indicated in each figure legend. Briefly, for comparisons of multiple groups over time or with two variables, a two-way analysis of variance (ANOVA) was used and corrected for multiple post hoc comparisons, comparing all groups to each other, all groups to a control, or selected groups to each other. For comparisons of multiple groups with only one variable, a one-way ANOVA was performed and corrected for multiple post hoc comparisons. For comparisons of two groups, two-tailed paired or unpaired t tests were used, except where indicated. Statistical analyses were performed in the GraphPad Prism 8 software. The non-parametric Kruskal–Wallis test with a Dunn’s post-hoc test was used to assess the difference in median abundance of eukaryotic viral and phage taxa in the Non-IBD, UC and CD cohorts. *P* values denoted throughout the manuscript highlight biologically relevant comparisons. A *P* value of less than 0.05 was considered significant, denoted as \**P* ≤ 0.05, \*\**P* ≤0.01, \*\*\**P* ≤ 0.001, and \*\*\*\**P* < 0.0001 for all analyses. FDR correction was performed for all RNA-seq data.

---

**Fig. S1.**
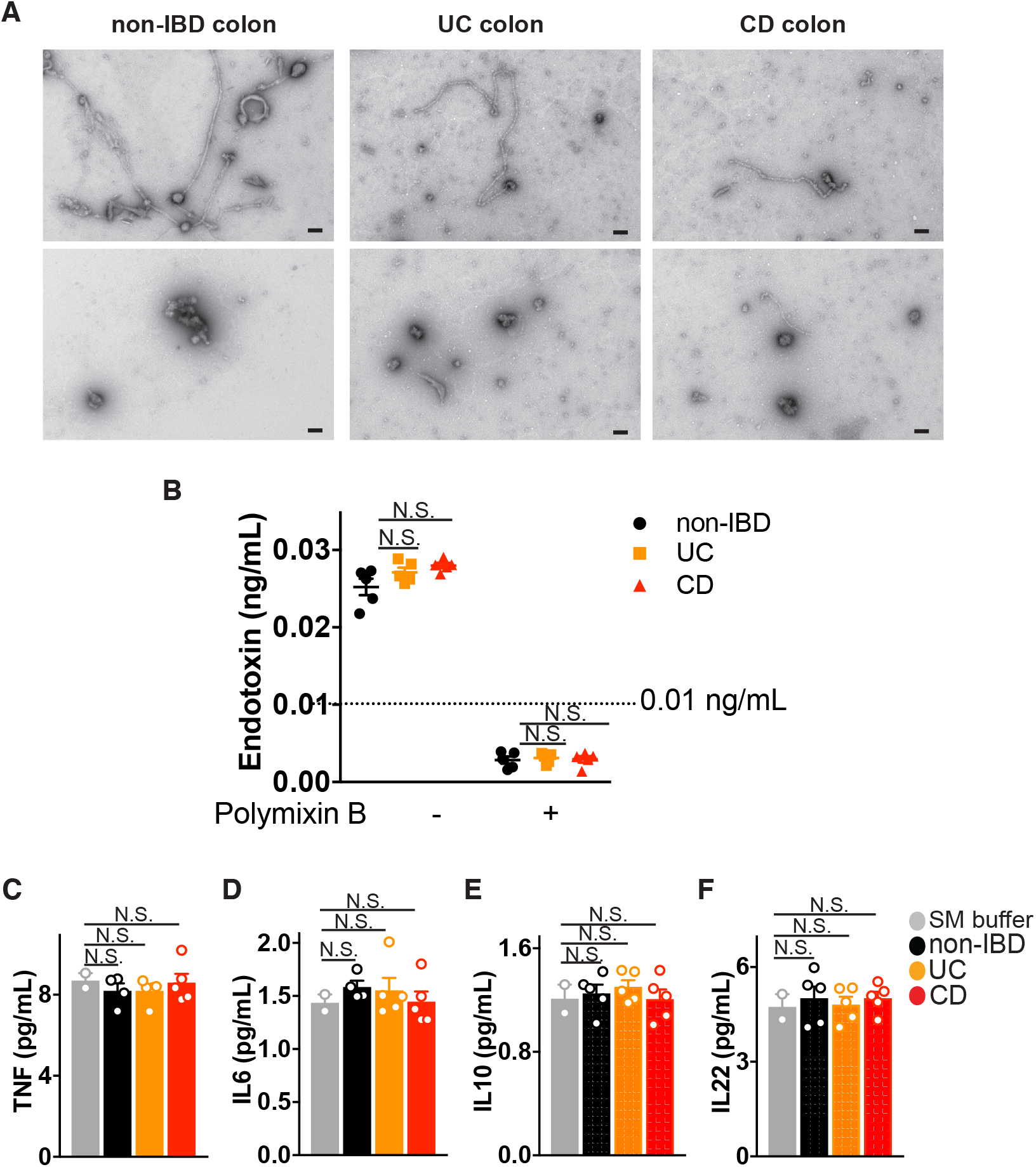
Isolation of virus-like particles from human colon resections. (A) Representative electron microscopy of virus-like particles (VLPs) isolated from fresh colon resection samples from non-IBD, Ulcerative Colitis (UC) or Crohn’s disease (CD) patients, post-surgery using uranyl acetate (scale bar, 100 nm). (B) Endotoxin levels in VLPs isolated from colon resections from non-IBD, UC or CD patients as measured by Limulus amebocyte lysate assay. (C) Tumor Necrosis Factor (TNF), (D) Interleukin (IL)-6, (E) IL-10 or (F) IL-22 levels in VLP preparations or buffer controls as measured by ELISA. Data is mean ± s.e.m. of 3-8 biological replicates, and one-way ANOVA with Tukey’s multiple comparison test was used.

**Fig. S2.**
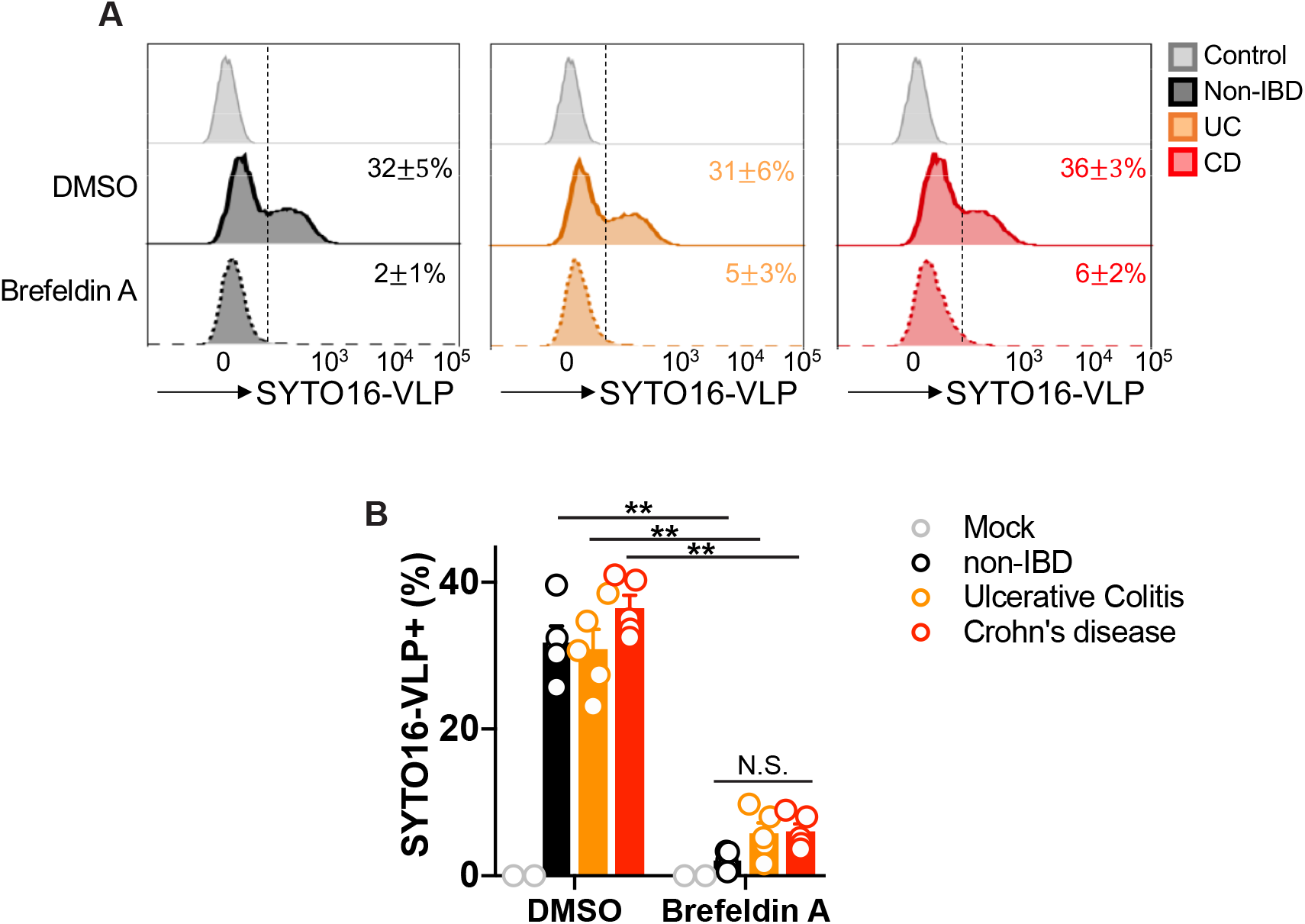
Internalization of intestine-resident viruses is endocytosis-dependent. (A) Flow cytometric analysis of SYTO-16 labeled virus like particles (VLPs) uptake by primary peripheral blood derived human macrophages 3 hours post incubation with 10 non-IBD, Ulcerative Colitis (UC) or Crohn’s disease (CD)-colon resection derived VLPs/cell. Uptake was assessed in the presence of Brefeldin A (3.0 mg/mL) or DMSO control. (B) Quantification of % of macrophages that took up SYTO-16 labeled VLPs from 5-8 individuals. Data is mean ± s.e.m. of 5-8 biological replicates. \*\**P*<0.01 and as determined by a one-way ANOVA with Tukey’s multiple comparison test.

**Fig. S3.**
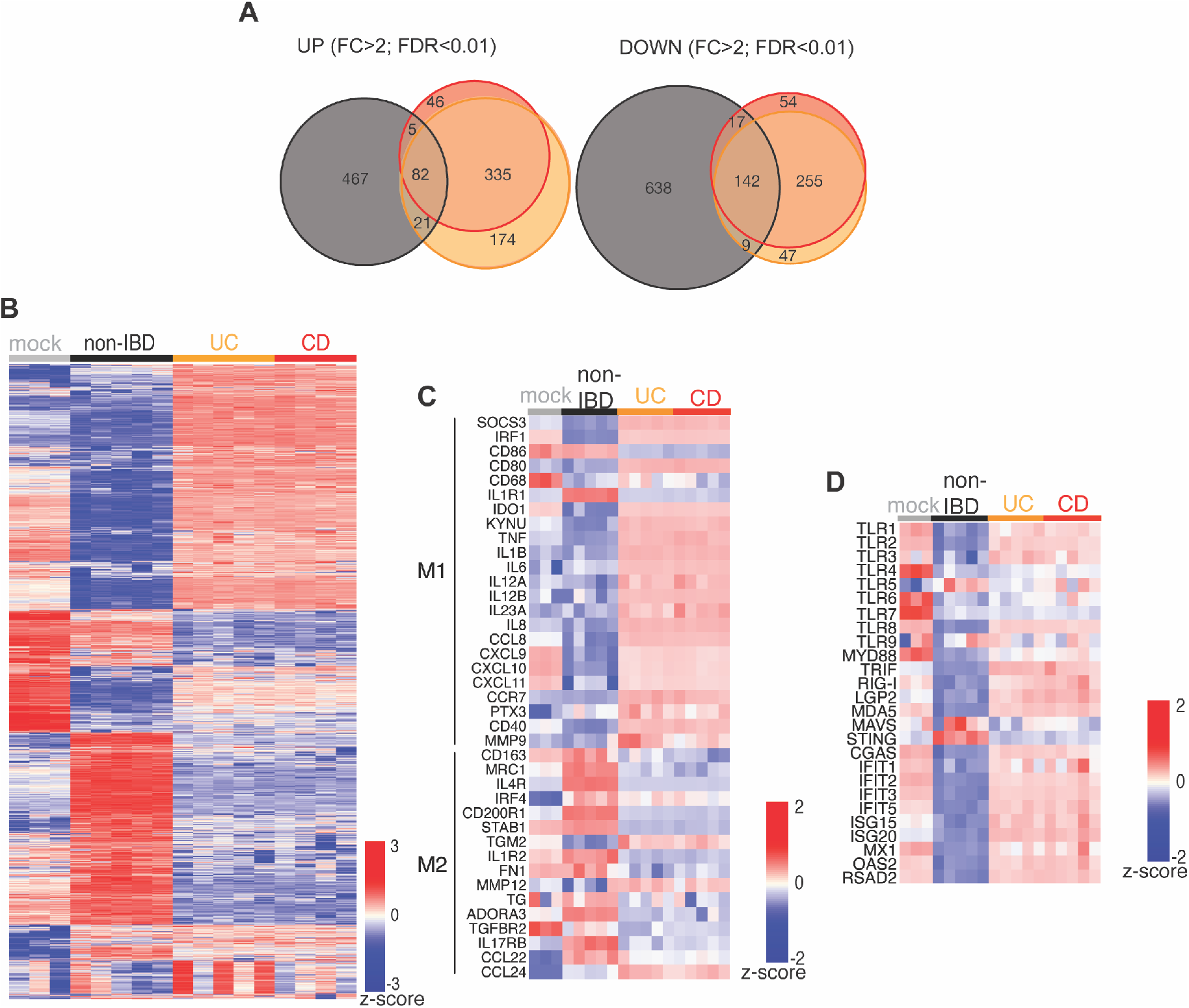
Divergent macrophage states induced by enteric viruses from non-IBD or IBD patient colons. (A) Venn diagram of genes significantly (Fold Change, FC>2; False Discovery Rate, FDR<0.01) up- or down-regulated by non-IBD, UC or CD VLPs. (B) Hierarchical clustered heatmap of z-score for differentially expressed genes (Fold change (FC) > 2, false discovery rate (FDR) < 0.05) following exposure of human peripheral blood derived macrophages to mock (SM buffer, n=3) or VLPs from non-IBD (n=5), Ulcerative Colitis (UC, n=5) or Crohn’s disease (CD, n=5) patient colon resections (10 VLPs/cell, 24h). (C) Heatmap of z-score for select genes defining “M1” pro-inflammatory or “M2” resolving macrophages or (D) Pattern Recognition Receptors (PRRs), signaling adapters and interferon stimulatory genes (ISGs).

**Fig. S4.**
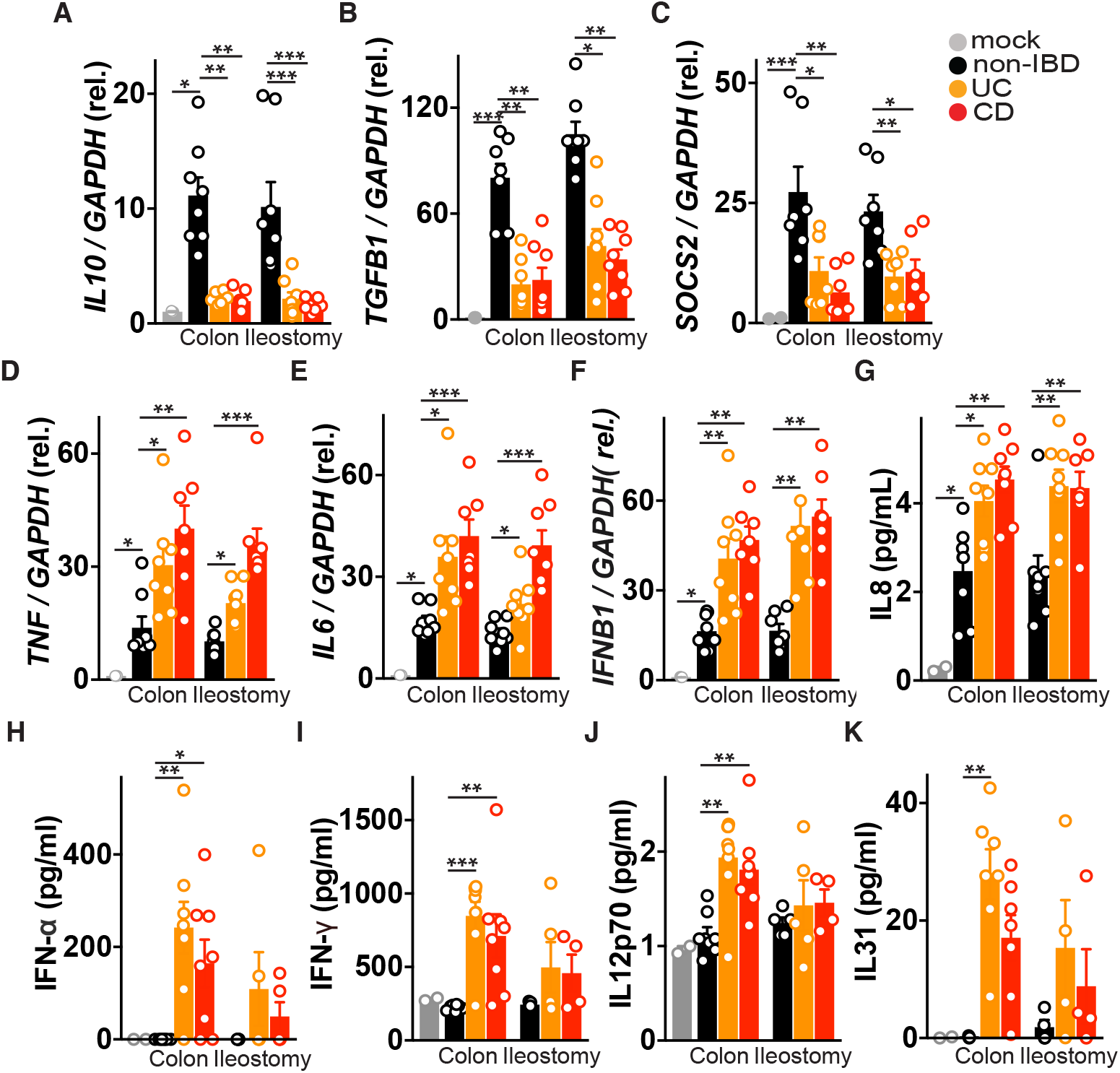
Enteric viruses from non-IBD patients protect while IBD viromes promote inflammation. Human peripheral blood derived macrophages were delivered virus-like particles (VLPs, 10 VLPs/cell, 24h) isolated from non-IBD controls, Ulcerative Colitis (UC) or Crohn’s disease (CD) patient colon resections or ileostomy fluid and (A) *IL10*, (B) *TGFB1*, (C) *SOCS2*, (D) *TNF*, (E) *IL6*, and (F) *IFNB1* mRNA levels were measured by qPCR. (G) IL-8, (H) IFN-*α*, (I) IFN-*γ*, (J) IL12p70 and (K) IL-31 production were measured by ELISA. Data is mean ± s.e.m. of 4-8 biological replicates. \**P* <0.05, \*\**P*<0.01, \*\*\**P*<0.001, as determined by a one-way ANOVA with Tukey’s multiple comparison test.

**Fig. S5.**
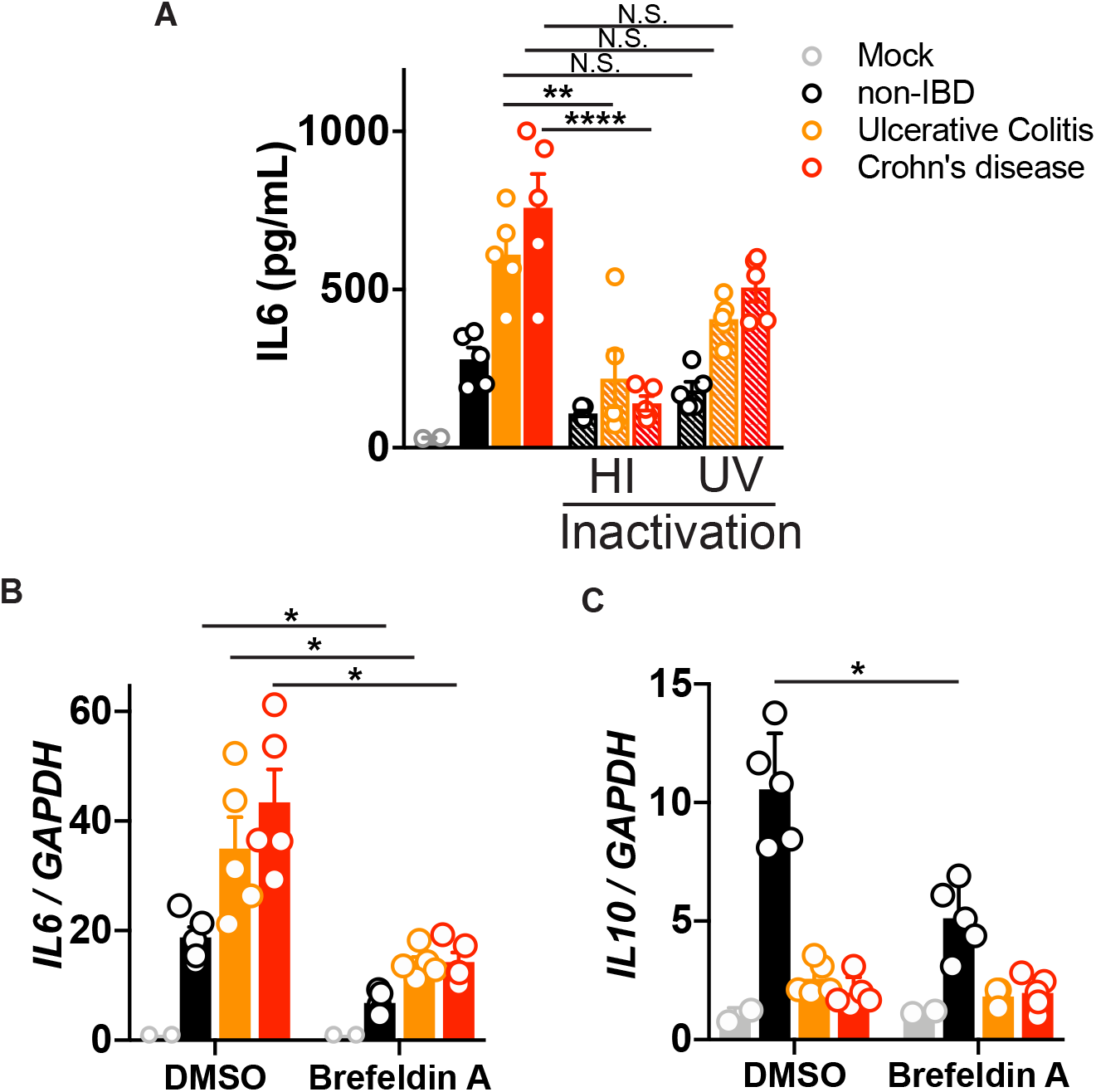
Requirement of endocytosis but not virus replication for virome immunodulation. (A) Interleukin (IL)-6 production from primary human peripheral blood-derived macrophages 24 hours following exposure to non-IBD, Ulcerative Colitis (UC) or Crohn’s disease (CD) colon resection-derived VLPs, heat-inactivated (60°C for 1h) VLPs, or UV-irradiated (three times 200 mJ/cm^2^ for 15 minutes) VLPs, as measured by ELISA. (B) IL-6 or (C) IL-10 levels in human peripheral blood-derived macrophages 24 hours following exposure to non-IBD, UC or CD colon resection-derived VLPs in the presence of Brefeldin A (3.0 mg/mL) or DMSO control. Data is mean ± s.e.m. of 5 biological replicates. \**P*<0.05, \*\**P*<0.01, \*\*\**P*<0.001, \*\*\*\**P*<0.0001, as determined by a one-way ANOVA with Tukey’s multiple comparison test.

**Fig. S6.**
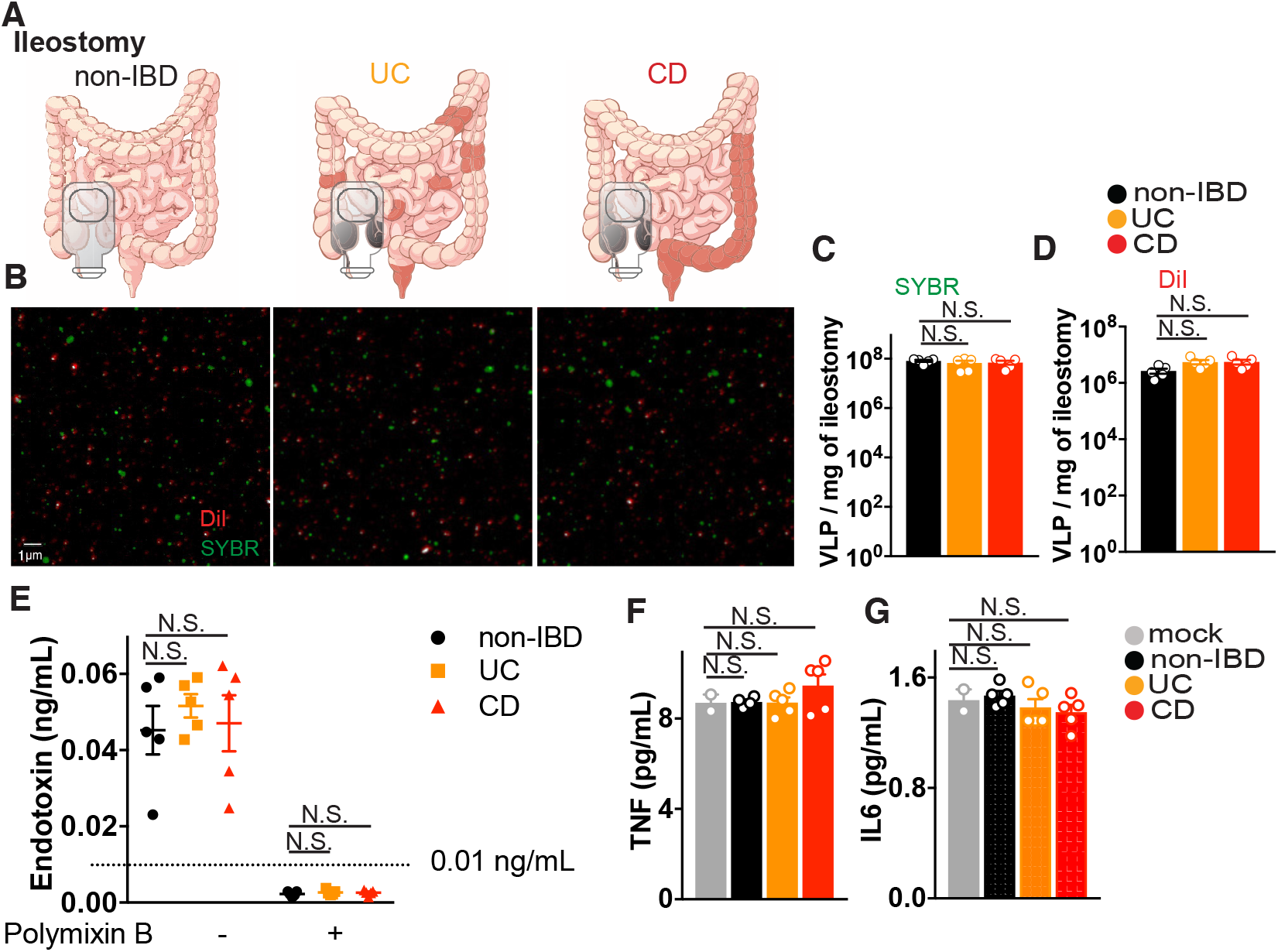
Isolation of virus-like particles from ileostomy pouch fluid. (A) Virus-like particles (VLPs) were isolated from ileostomy fluid collected from non-IBD, Ulcerative colitis (UC) or Crohn’s disease (CD) patients (Table S2). (B) Confocal imaging of VLPs isolated from ileostomy fluid of non-IBD, UC or CD patients (C and D) Quantification of viral nucleic acid using SYBR Gold (C) and phospholipid bilayers using Dialkylcarbocyanine Probe, DiI (D). (E) Endotoxin levels in VLP preps isolated from ileostomy contents of non-IBD, UC or CD patients measured by Limulus amebocyte lysate assay. (F) TNF and (G) IL-6 levels in VLP preps from ileostomy fluids as measured by ELISA. Data is mean ± s.e.m. of 5-8 biological replicates. Graphs are representative of 2-3 experiments and n ≥5 replicates, and one-way ANOVA with Tukey’s multiple comparison test was used.

**Fig. S7.**
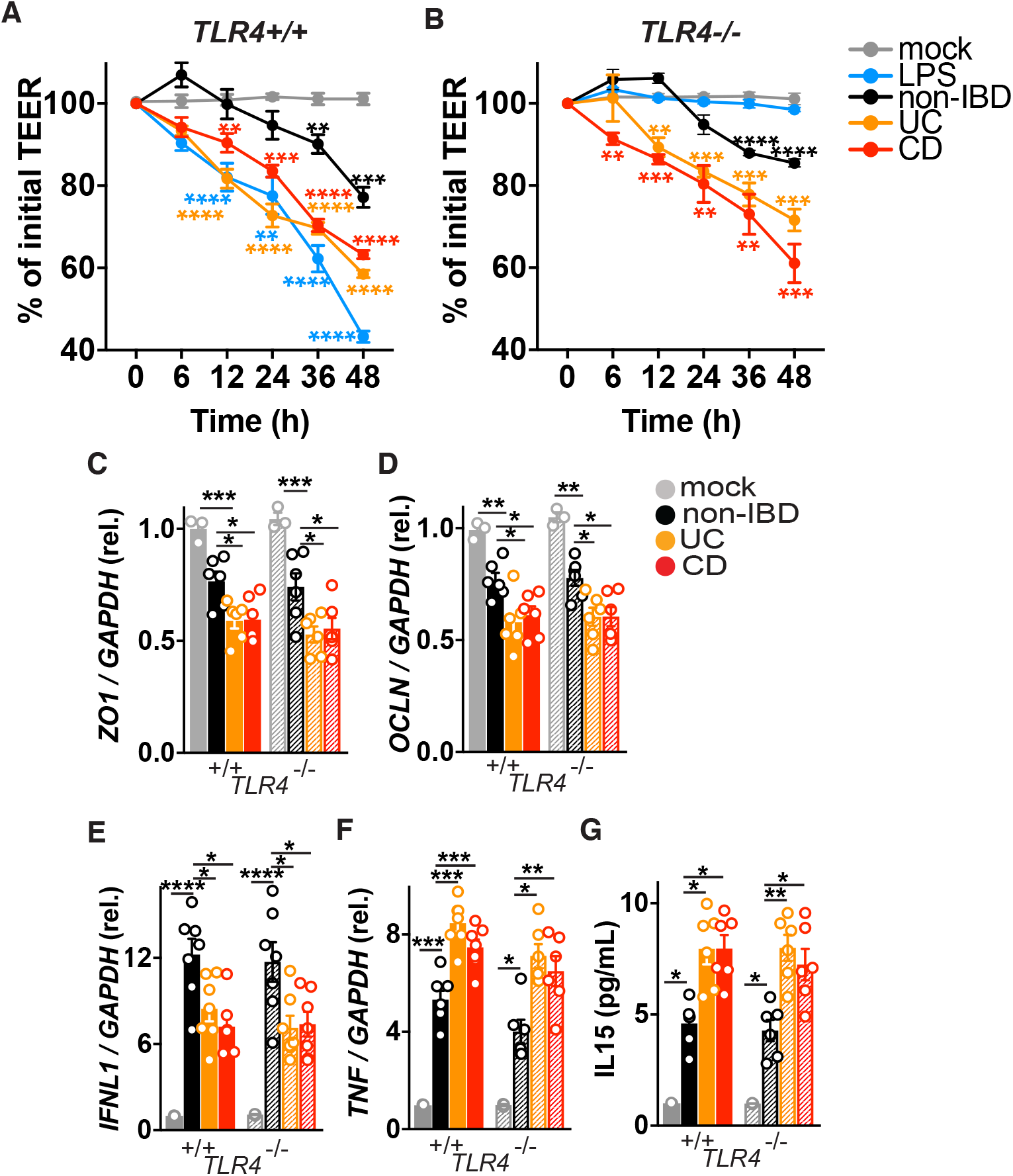
IBD enteric viruses disrupt intestinal epithelial barrier integrity in a TLR4-independent manner. (A) Trans-epithelial electrical resistance (TEER) over time of wild-type or (B) TLR4^−/−^ Caco2 epithelial monolayers following stimulation with LPS (100 ng/mL) or exposure to SM buffer (mock), non-IBD, Ulcerative colitis (UC) or Crohn’s disease (CD) colon resection-derived VLPs (10 VLPs/cell). (C) Zonula occludens (*ZO1*), (D) *Occludin (OCLN),* (E) *IFNL1*, (F) *TNF* transcript expression or (G) IL-15 levels 48h post-incubation of monolayer wild-type or TLR4 knockout Caco2 cells with VLPs isolated from non-IBD, UC and CD colon resections as measured by qPCR or ELISA. Data is mean ± s.e.m. of 5-8 biological replicates. \**P* < 0.05, \*\**P*<0.01, \*\*\**P*<0.001, as determined by a one-way ANOVA with Tukey’s multiple comparison test.

**Fig. 8.**
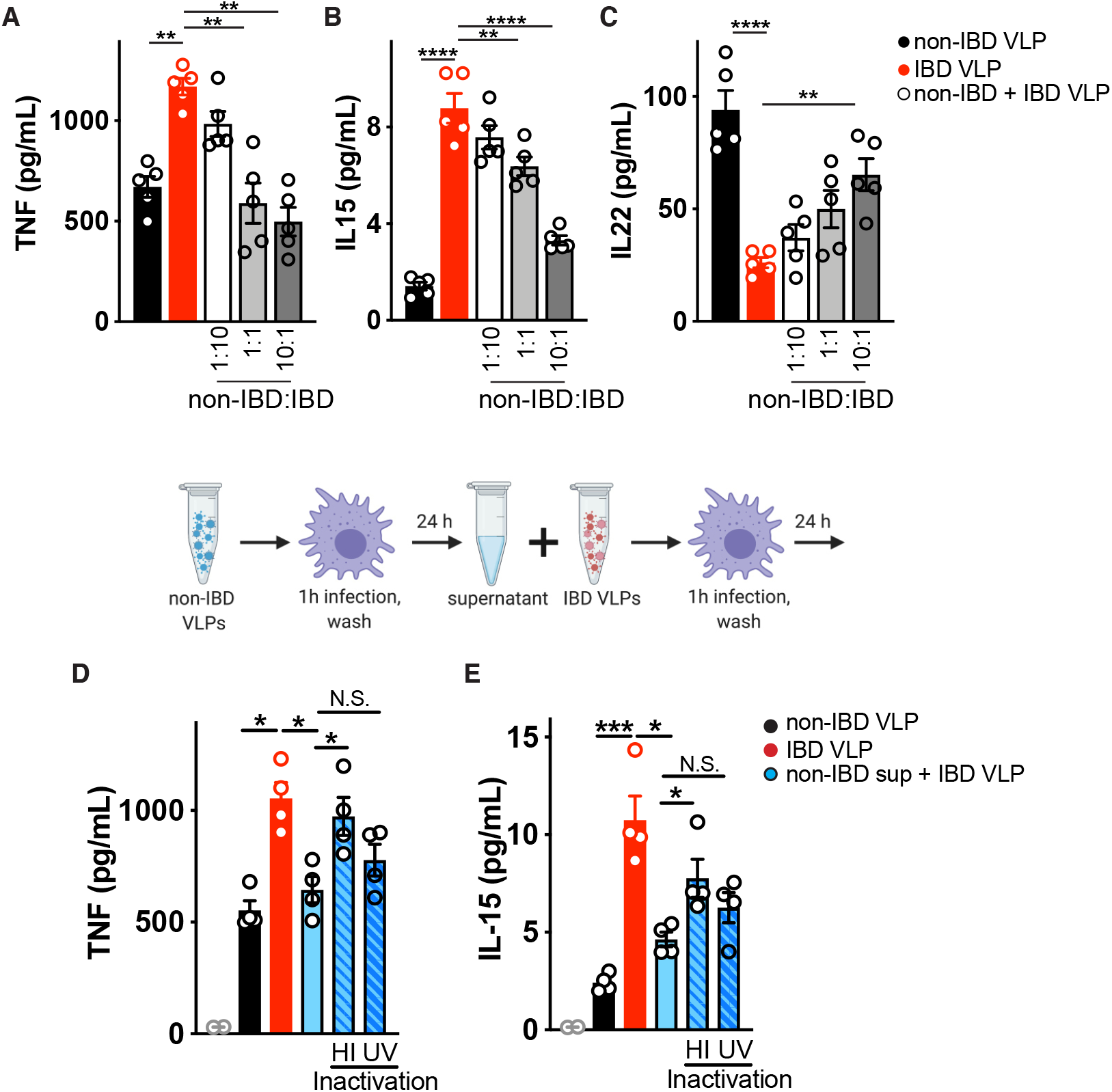
Suppression of IBD virome-induced inflammation by healthy human enteric viromes. Primary human peripheral blood-derived macrophages were delivered colon resection-derived non-IBD VLPs (10 VLPs/cell), IBD VLPs (10 VLPs/cell) or indicated ratios of non IBD to IBD VLPs and (**A**) TNF, (**B**) IL-15, and (**C**) IL-22 levels were measured at 24h by ELISA. Schematic for incubation of human primary macrophages with IBD VLPs and 24h supernatant from non-IBD VLP exposed macrophages. (**D**) TNF, and (**E**) IL-15 levels measured by ELISA following IBD VLP delivery to macrophages with the addition of non-IBD macrophage supernatant that was heat inactivated (95°C for 10 min) or UV crosslinked (three times 200 mJ/cm^2^ for 15 minutes) as controls. Data are mean ± s.e.m. of 5 biological replicates. \**P*<0.05, \*\**P*<0.01, \*\*\**P*<0.001, \*\*\*\**P*<0.0001 as determined by one-way ANOVA with Tukey’s multiple comparison test.

**Fig. S9.**
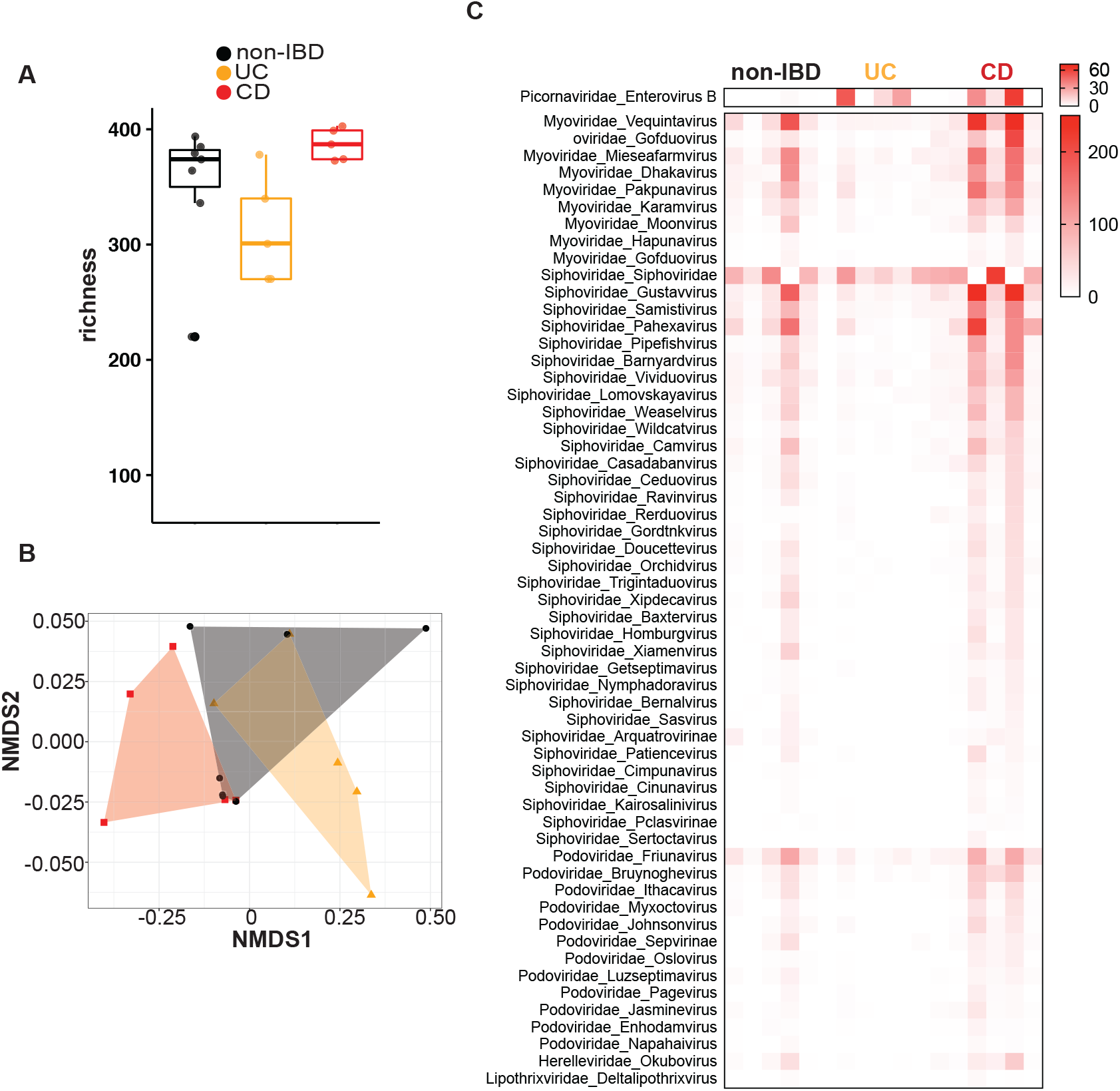
Characterization of VLPs in colon resections from non-IBD, Ulcerative Colitis and Crohn’s disease patients. VLP were assessed using VLP DNA and RNA metagenomic sequence data. Sequences were assigned a eukaryotic viral or phage taxonomic lineage using primary searches against virus sequence databases and subsequently confirmed in secondary searches using reference databases with additional non-viral taxonomic lineages. Specific eukaryotic viral families were selected for subsequent analysis by gating on the alignment length and percent identity for each family that separated the high-quality alignments from potentially spurious alignments (Table S3). (A) VLP species richness was calculated using the spec number function of the Vegan package. Significance was assessed via Kruskal–Wallis test with a Dunn’s post-hoc test. (B) Non-metric multidimensional scaling (NMDS) using Bray-Curtis dissimilarity was conducted using the metaMDS function in Vegan and PERMANOVA analysis (permutational multivariate analysis of variance) was performed using the pairwise Adonis package to assess significance. There were no significant differences in beta diversity between the groups. (C) Significantly (P-adjusted < 0.05, Kruskal–Wallis test with a Dunn’s post-hoc correction) differentially abundant eukaryotic and prokaryotic viruses in UC or CD colon resections. Data are the mean of 5-7 biological replicates.

**Fig. S10.**
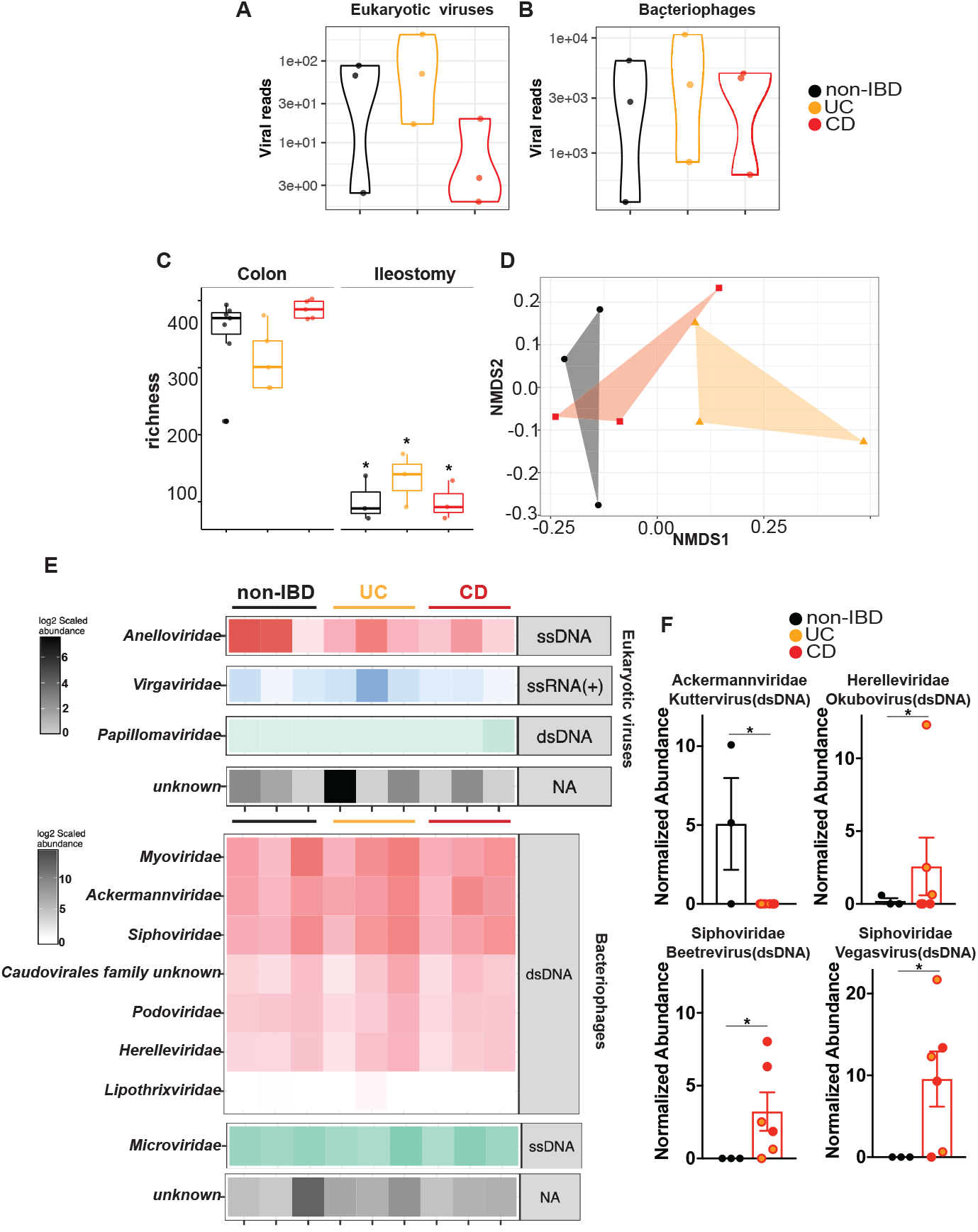
Detection and characterization of VLPs in ileostomy fluid from non-IBD, Ulcerative Colitis and Crohn’s disease patients. (A, B) Violin plots represent the distribution of individual datasets for total reads aligning to eukaryotic viruses or bacteriophages from non-IBD, UC or CD cohorts. (C) Phage VLP richness was assessed using VLP metagenomic sequence data and the spec number function of the Vegan package. Significance was assessed via Kruskal–Wallis test with a Dunn’s post-hoc test. (D) NMDS using Bray-Curtis dissimilarity was conducted using the metaMDS function in Vegan and PERMANOVA analysis was performed using the pairwise Adonis package to assess significance. There were no significant differences in beta diversity between the groups. (E) Taxonomic assignments of VLP sequences with high quality alignments in non-IBD, UC or CD ileostomy samples. The families are grouped by their Baltimore Classification and ranked by relative abundance in the dataset. (F) Significantly differentially abundant genera of bacteriophages in ileostomy samples from non-IBD versus IBD cohorts. Data are the mean of 5-7 biological replicates. \**P*<0.05, Kruskal–Wallis test with a Dunn’s post-hoc test.

**Fig. S11.**
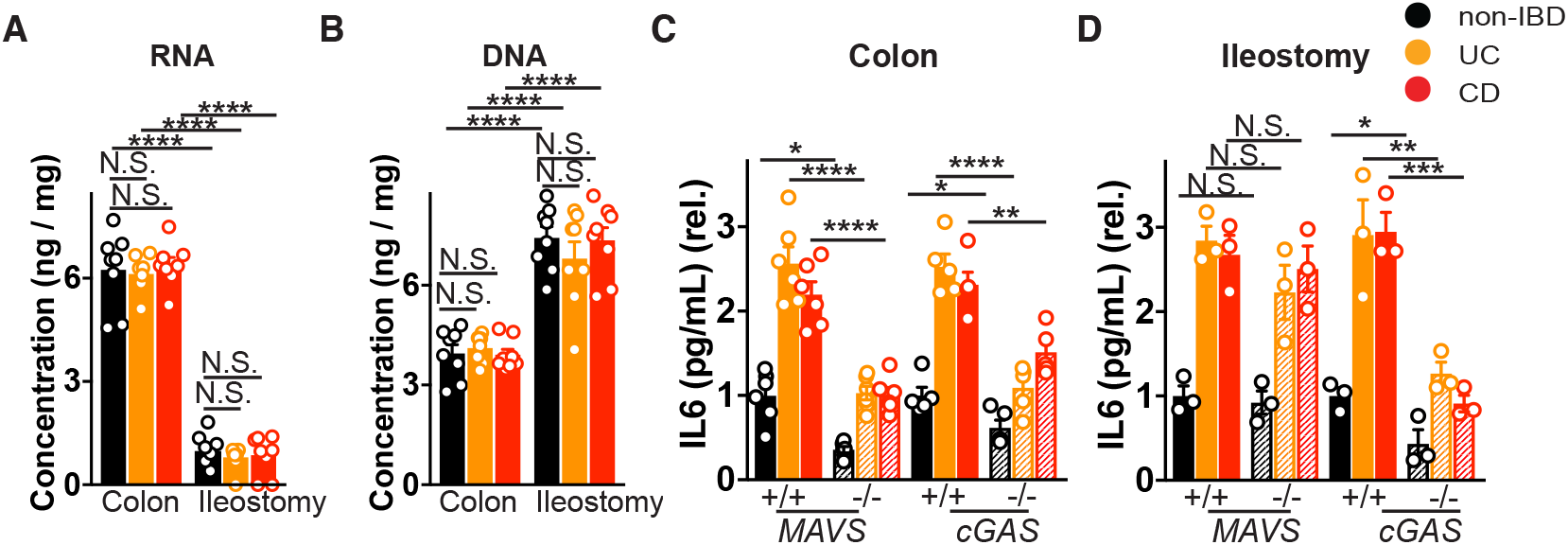
Differential host receptor requirement for immune response to VLPs isolated from colon or ileostomy fluid. Concentration of (A) RNA, and (B) DNA isolated from non-IBD, Ulcerative colitis (UC) or Crohn’s disease (CD) colon resections or ileostomy fluid derived VLPs. (C) IL-6 levels following exposure of bone marrow derived macrophages (BMDMs) from wild-type, MAVS^−/−^ or cGAS^−/−^ mice to VLPs (10 VLPs/cell, 24h) derived from (C) non-IBD, UC or CD colon resections or (D) ileostomy fluid as measured by ELISA. Data is mean ± s.e.m. of 3-6 biological replicates. \**P*<0.05, \*\**P*<0.01, \*\*\**P*<0.001, \*\*\*\**P*<0.0001, as determined by a one-way ANOVA with Tukey’s multiple comparison test.

**Fig. S12.**
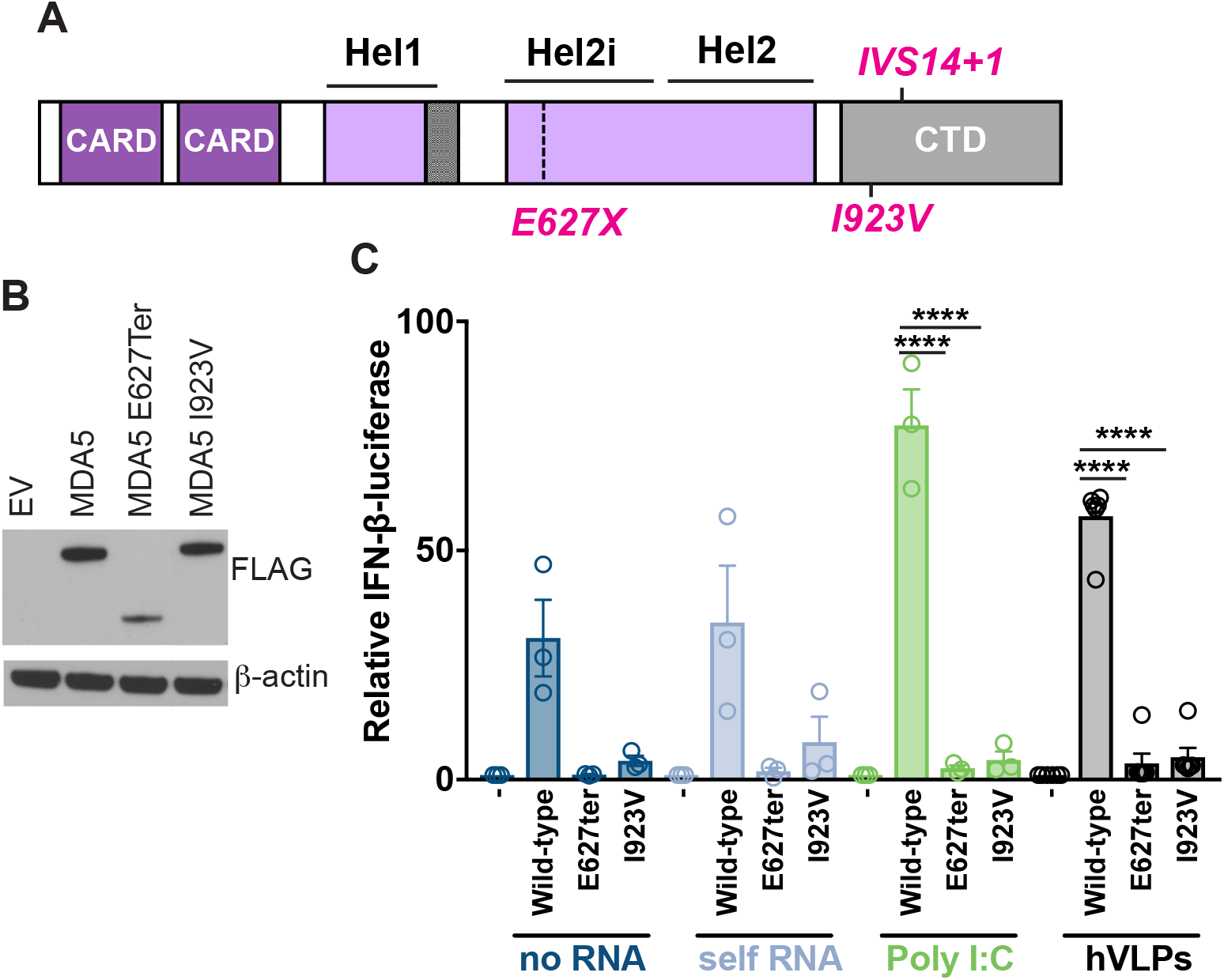
IBD-risk loss-of-function MDA5 exhibits dampened responses to the human enteric virome. (A) Schematic of human MDA5 and mapped IBD-associated MDA5 variants rs35744605/E627X, rs35667974/I923V and rs35732034/IVS14+1. (B) Western Blot of ectopically expressed, FLAG-tagged MDA5 in HEK293T cells. (C) Relative activation of IFNβ luciferase reporter in HEK293T cells transfected with Empty vector (EV), FLAG-tagged WT MDA5, MDA5E627Ter or MDA5I923V and transfected with 1μg/mL RNA isolated from the cytoplasm of HEK293T cells (self RNA), poly I:C or RNA isolated from human colon resection virus like particles (VLPs) for 24 hours.

**Fig. S13.**
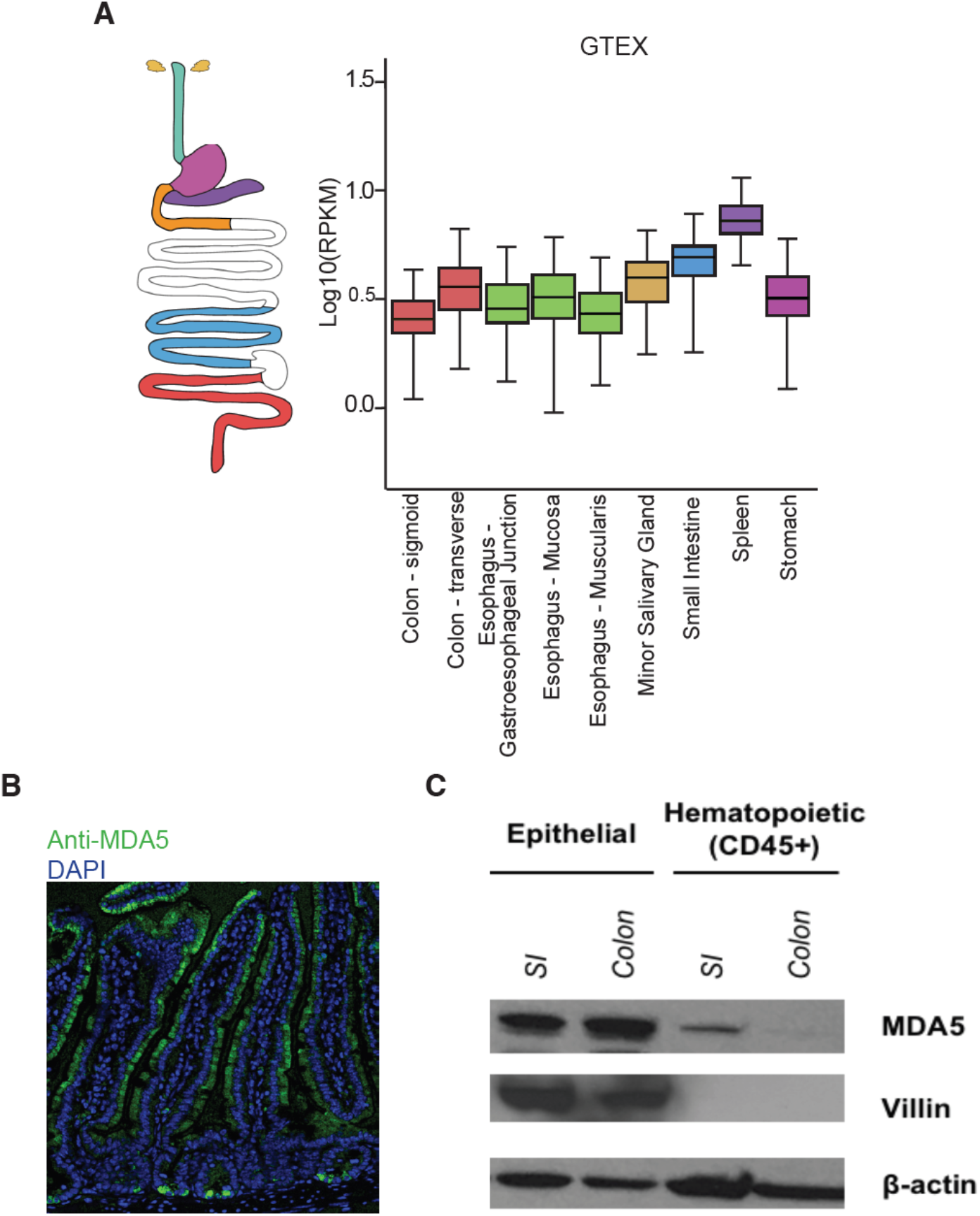
Virus receptor MDA5 is preferentially expressed in intestinal epithelium. (A) *IFIH1* (MDA5) mRNA levels across several gastrointestinal-related human tissues and spleen were retrieved from Genotype-Tissue Expression (GTEX) Portal (Broad Institute) (B) Confocal microscopy of MDA-5 (green) or nuclei (DAPI, blue) in mouse small intestine. (C) Immunoblot of MDA5 in epithelial cells or CD45+ hematopoietic cells sorted from mouse small intestine (SI) or colon.

**Fig. S14.**
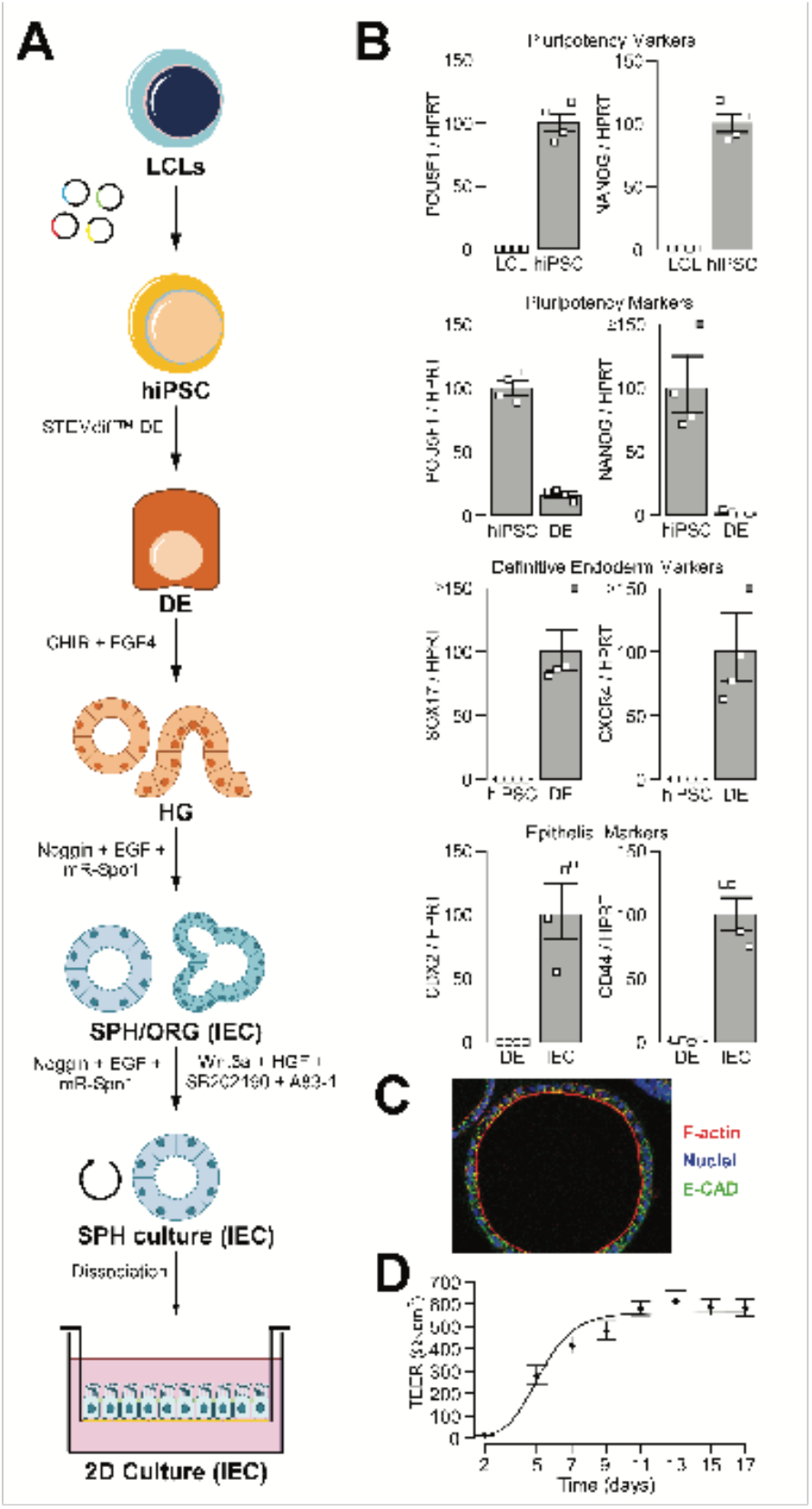
Generation of hiPSC-derived human intestinal epithelial cells. (A) Schematic illustrating the different steps involved in the generation of hiPSC-derived 2D epithelial cultures from LCLs. (B) Progression through the different steps of the differentiation protocol is measured via the expression of specific markers by qPCR: (1) conversion of LCLs to hiPSC is shown by the increased expression of pluripotency markers such as POU5F1 and NANOG; (2) differentiation of hiPSC to DE is confirmed through the loss of these pluripotency markers as well as with the gain of DE-specific markers such as SOX17 and CXCR4; (3) the generation of IEC cultures is validated by the expression of intestinal-specific markers such as CDX2 and CD44. (C) Epithelial cultures are also evaluated through confocal immunofluorescence imaging to confirm the formation of epithelial-like polarized structure (spheroids are shown; for IF of 2D epithelial monolayer see Fig 10): nuclei (DAPI, blue), F-actin (phalloidin, red), and E-cadherin (green). (D) Barrier integrity of epithelial monolayers is assessed by measurement of trans-epithelial electric resistance (TEER).

**Fig. S15.**
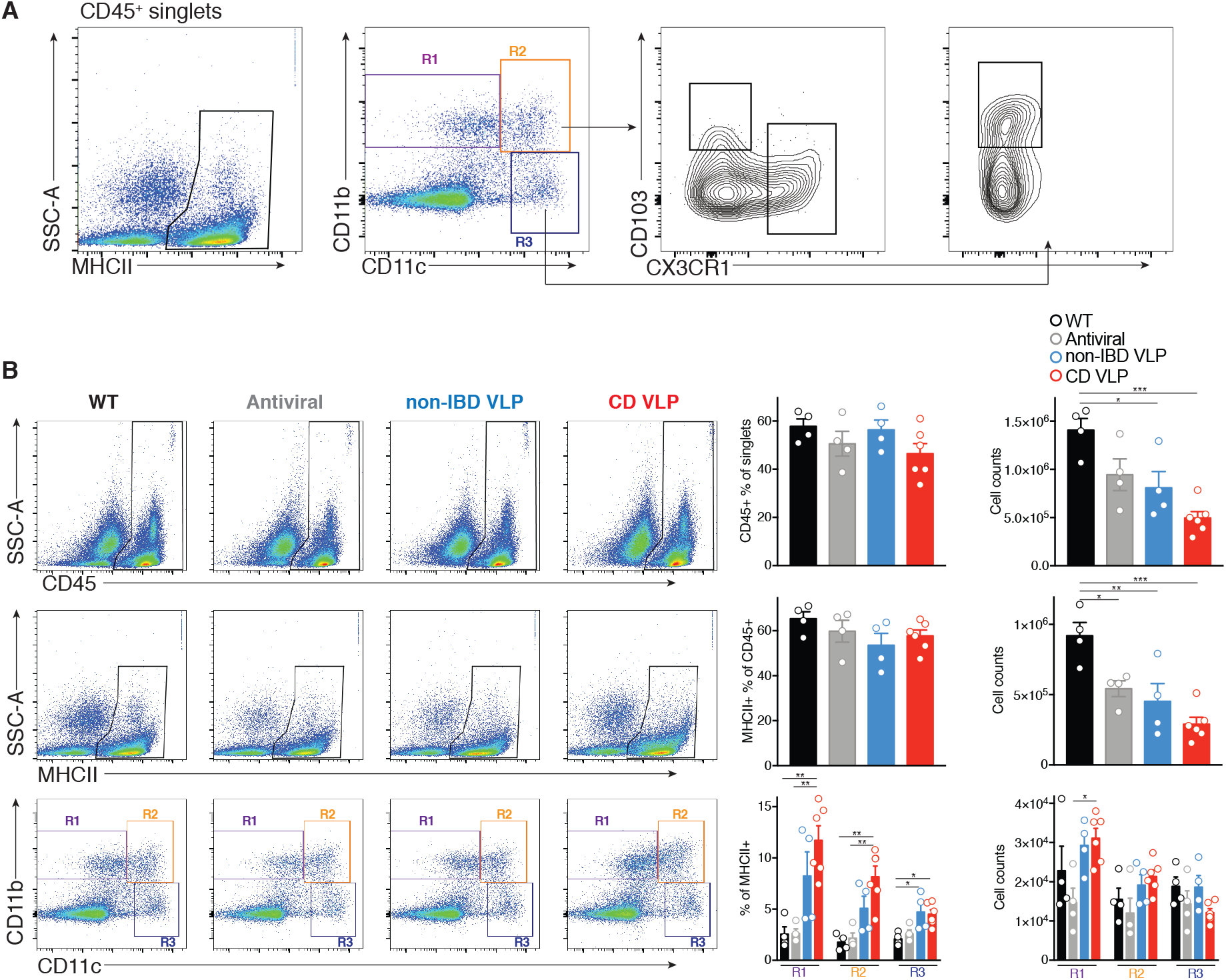
Quantification of mononuclear phagocytes in colonic lamina propria of humanized virome mice. C57BL/6 mice were depleted of their enteric virome by an antiviral cocktail (Acyclovir 20mg/kg, Lamivudine 10mg/kg, Ribavirin 30mg/kg, Oseltamivir 10mg/kg, gray, daily gavage for 10 days) and reconstituted with VLPs isolated from colonic resections of 4 non-IBD or 6 Crohn’s disease (CD) patients. (A) Colonic lamina propria were isolated as described in the methods section and analyzed using the described gating strategy. Cells were first gated on CD45+/MHCII+ live singlets and CD11b+CD11c- (R1), CD11b+CD11c+ (R2) and CD11b-CD11c+ (R3) mononuclear phagocytes were examined for expression of CX3CR1 or CD103. (B) Representative flow cytometry plots and quantification of frequency and the total count for the given immune cell subsets, including total CD45+ singlets (top), CD45+/MHCII+ singlets (middle), and CD11b or CD11c expressing CD45+/MHCII+ singlets (bottom). Each circle represents the cells from one mouse. Data is mean ± s.e.m. of 4-6 mice per group. \**P*<0.05, \*\**P*<0.01 and \*\*\**P*<0.001 as determined by a one-way ANOVA with Tukey’s multiple comparison test.

**Fig. S16.**
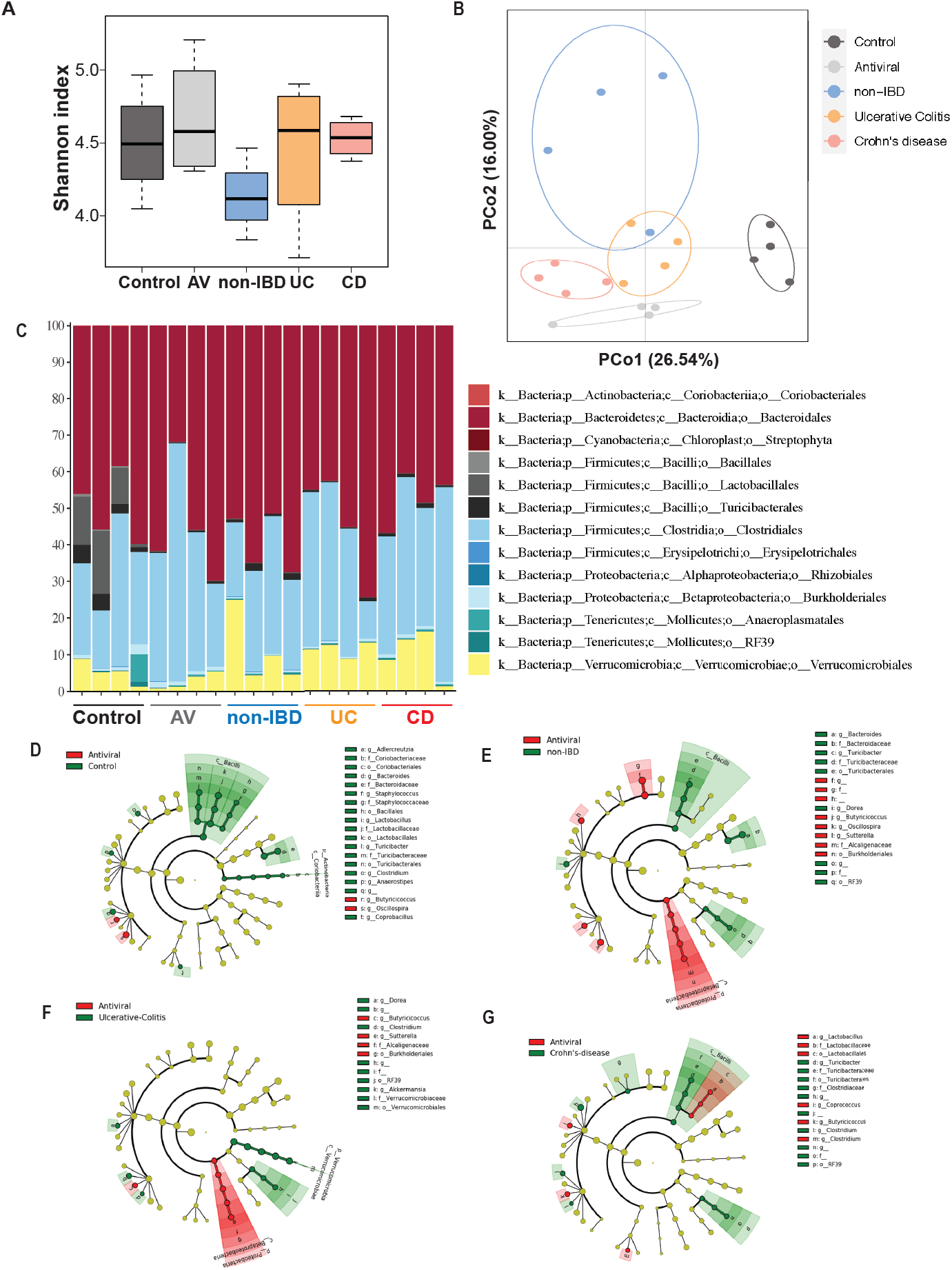
Abundance and composition of C57BL/6 murine bacterial communities following administration of human enteric viromes. Murine fecal bacterial communities before and after antiviral cocktail treatment and following gavage administration of healthy, Ulcerative colitis (UC) or Crohn’s disease (CD) human colon resection-derived viromes were determined using 16S rRNA sequencing. (A) Diversity of bacterial communities as determined by Shannon Index. (B) Principle Coordinate Analysis (PCoA), P < 0.05, PERMANOVA on unweighted UniFrac distance. (C) Distribution of operational taxonomic units (key) in feces; each operational taxonomic unit (selected at a sequence identity of 99%; QIIME software) is presented as relative abundance in each sample. (D) The compositional differences between control and antiviral treated mice, (E) antiviral treated compared to non-IBD virome humanized mice, (F) antiviral treated compared to UC virome humanized mice and (G) antiviral treated compared to CD virome humanized mice were determined by a linear discriminant analysis using LEfSe with a threshold of 2.0 on the logarithmic score using Galaxy (https://huttenhower.sph.harvard.edu/galaxy/).

**Table S1.**
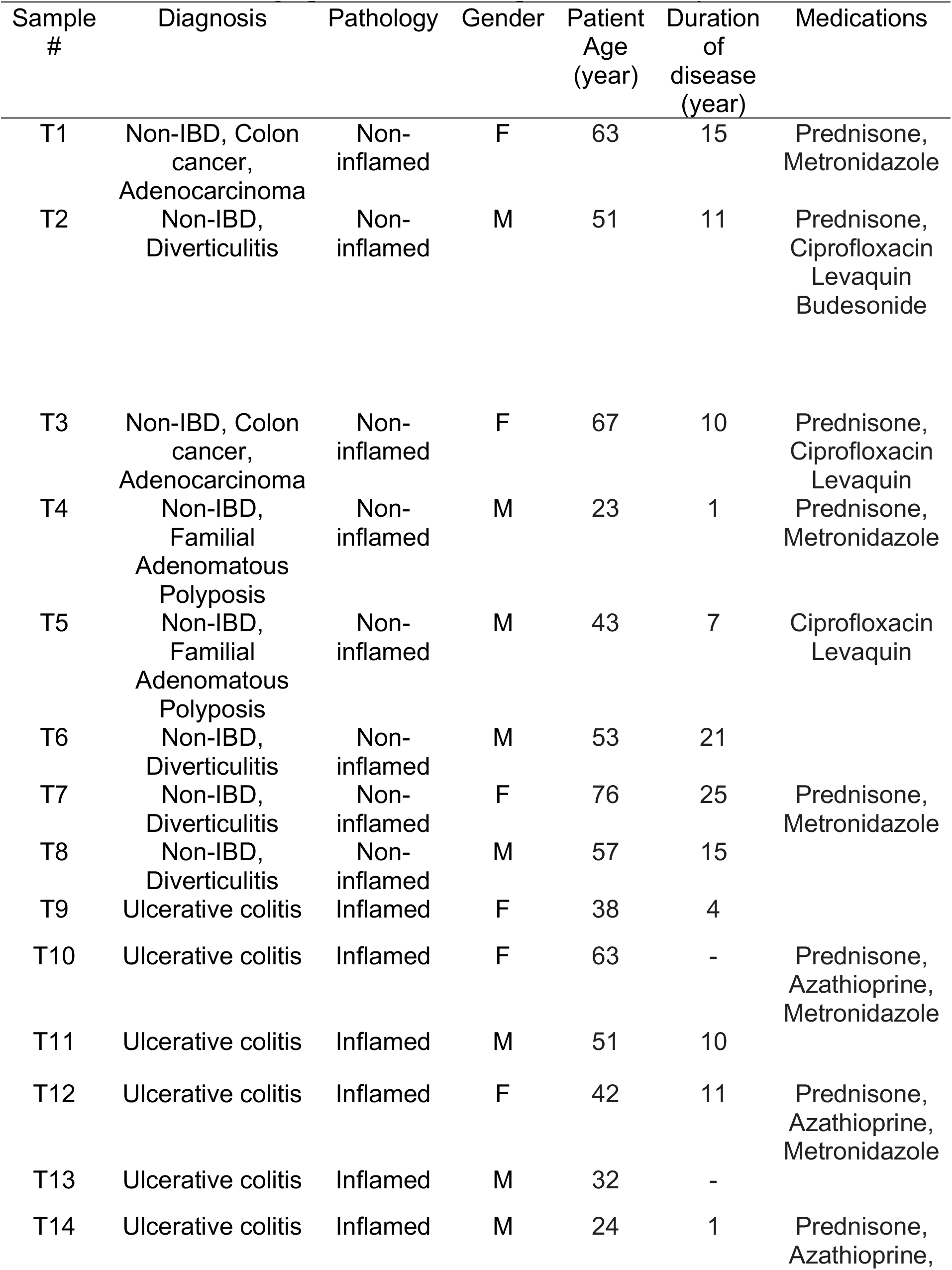
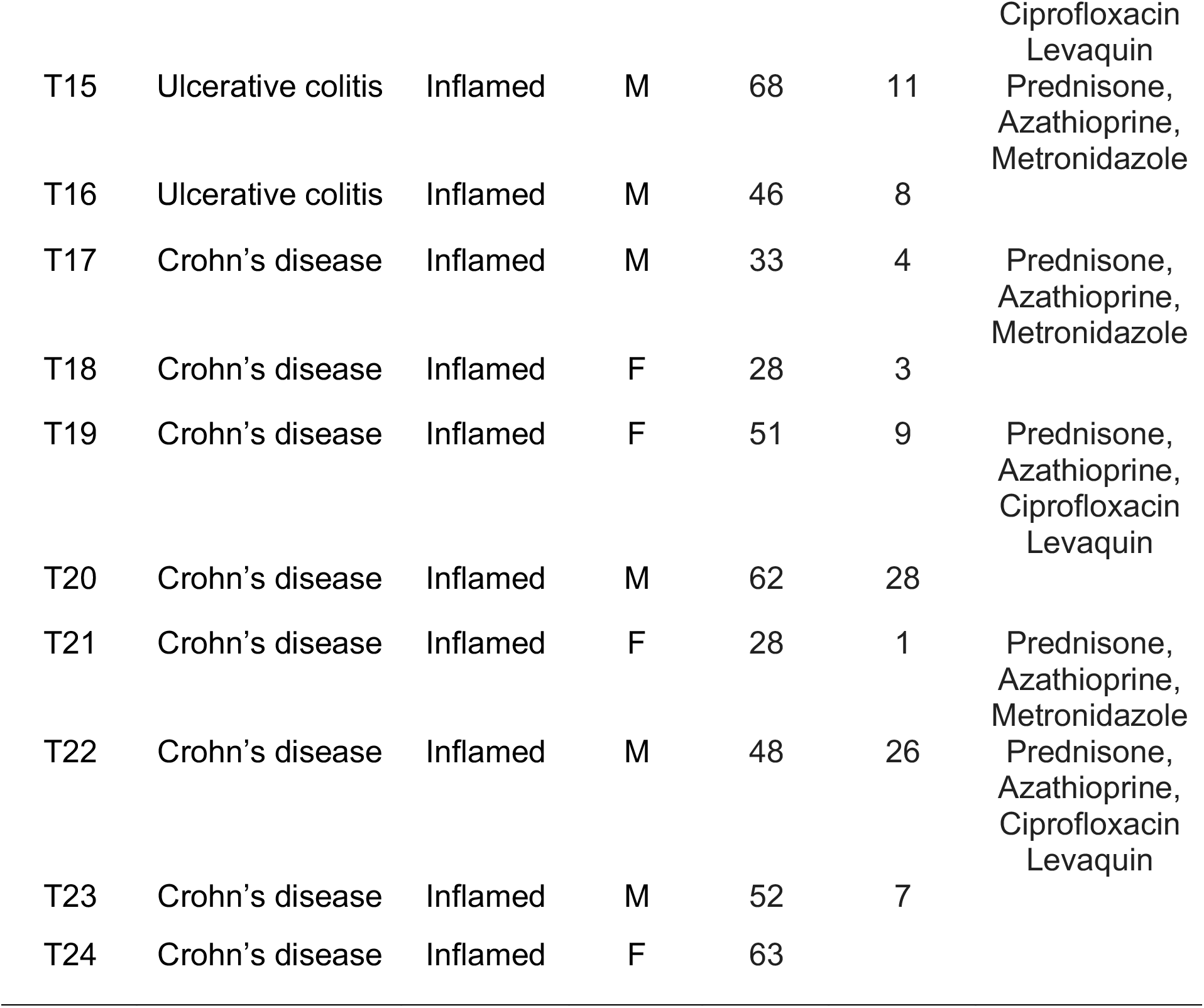
Demographic Data of Participants in the Study, Colon Resections.

**Table S2.**
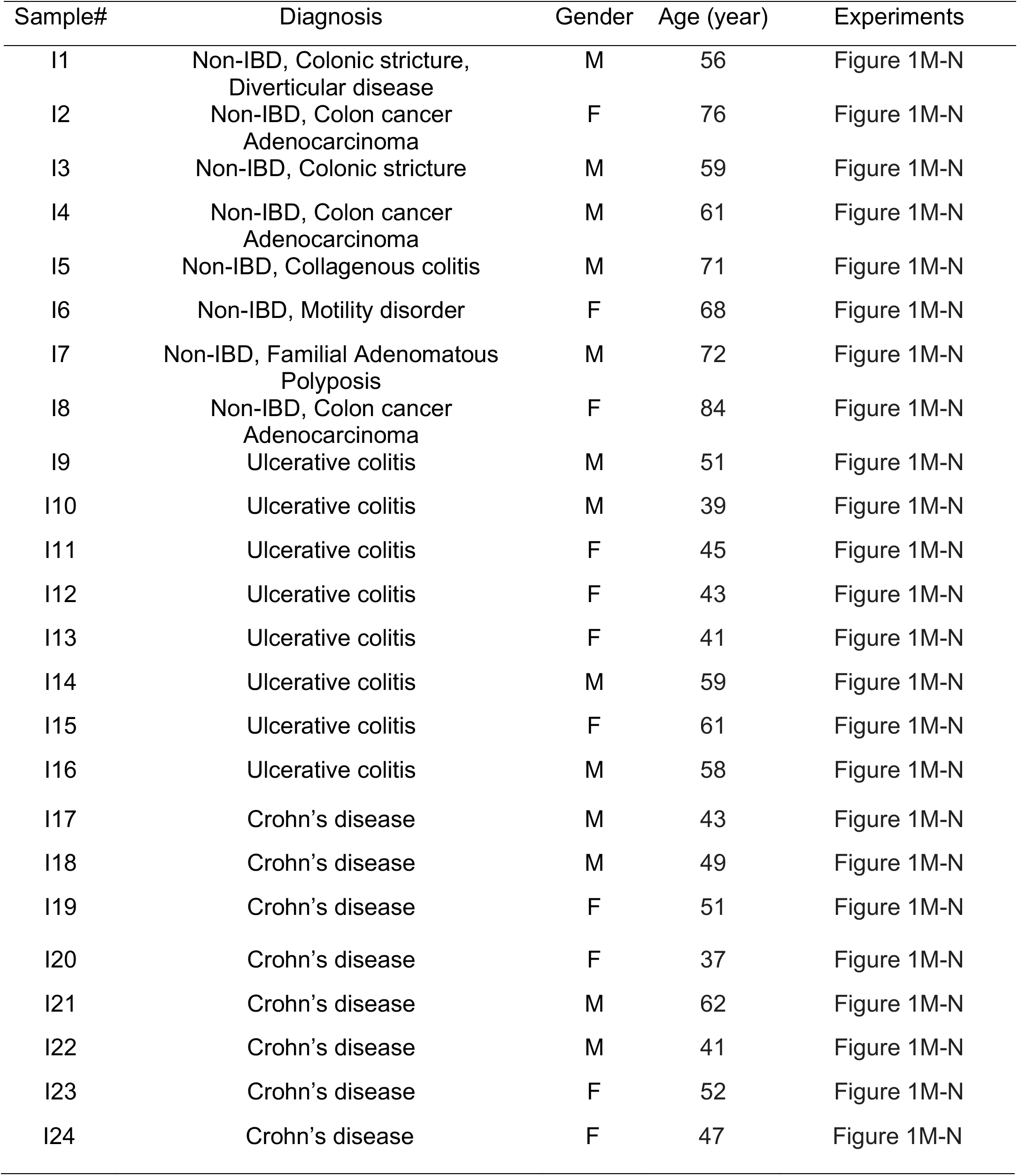
Demographic Data of Participants in the Study, Ileotomy Samples.

**Table S3.** Alignment length and percent identity requirements for confidently calling eukaryotic virus families.

| **Family** | **Minimum Percent ID** | **Minimum Alignment length in nt** |
| --- | --- | --- |
| **Herpesviridae** | **70** | **80** |
| **Picornaviridae** | **80** | **50** |
| **Papillomaviridae** | **70** | **75** |
| **Coronaviridae** | **70** | **100** |
| **Rhabdoviridae** | **70** | **75** |
| **Orthomyxoviridae** | **70** | **80** |
| **Anelloviridae** | **70** | **75** |
| **Virgaviridae** | **75** | **80** |

**Table S4.**
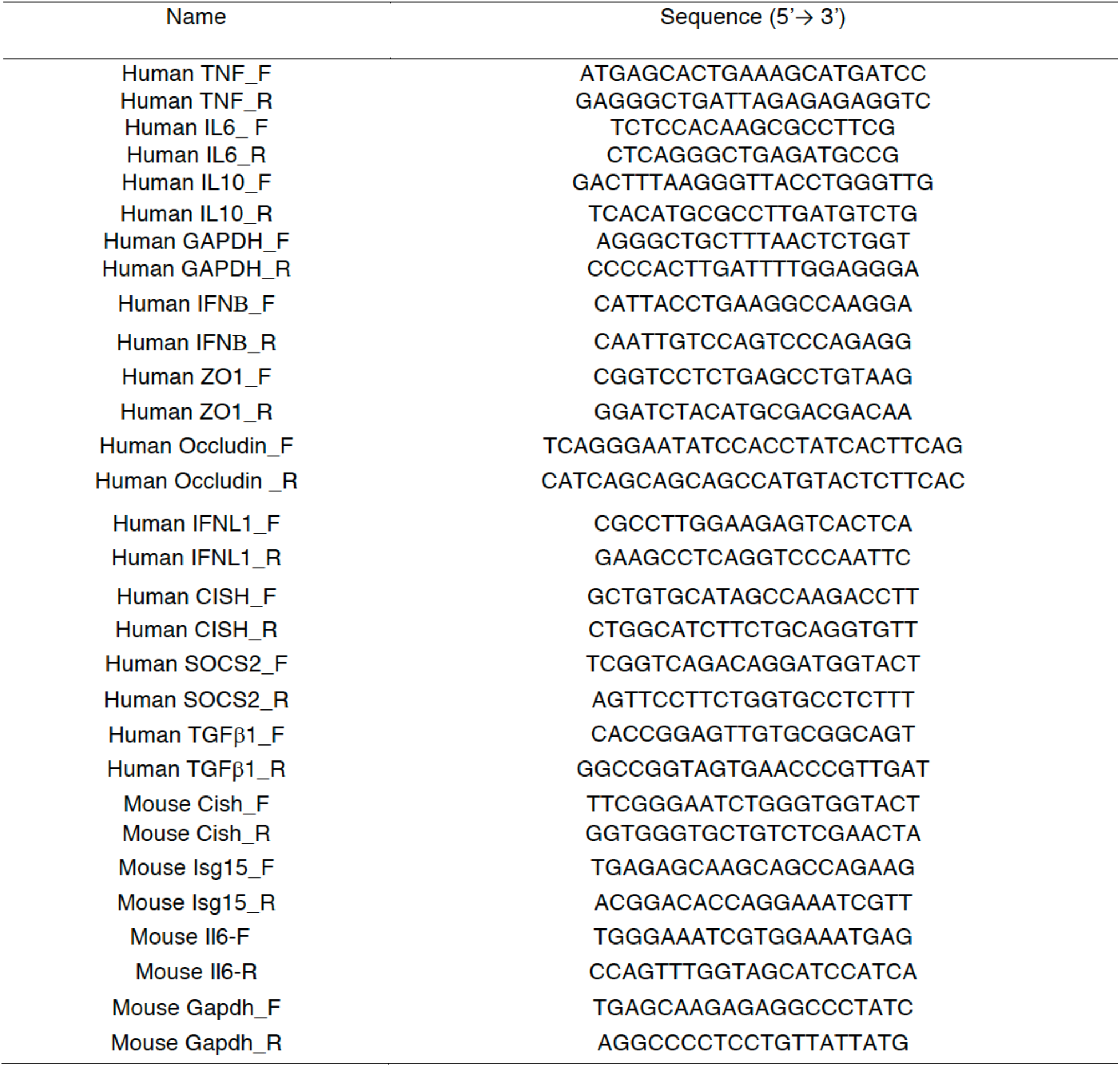
qPCR primer sequences.

**Table S5.** Pertinent information for commercial antibody reagents used for FACS analysis.

| Antibody | Supplier | Species | Cat # |
| --- | --- | --- | --- |
| CD45 (30-F11; Pacific Blue) | Biolegend | Mouse | 103101 |
| CD11b (clone M1/70; APC/Cy7) | Biolegend | Mouse | 101225 |
| CD11c (N418; PE) | Biolegend | Mouse | 117307 |
| Gr-1 (RB6-8C5; APC) | Biolegend | Mouse | 108411 |
| CX3CR1 (SA011F11; Brilliant Violet 785) | Biolegend | Mouse | 149029 |
| F4/80 (BM8; Brilliant Violet 650) | Biolegend | Mouse | 123149 |
| CD103 (2E7; PerCP/Cy5.5) | Biolegend | Mouse | 121415 |
| MHCII (M5/114.15.2; Alexa Fluor 700) | Biolegend | Mouse | 107621 |

## Notes

### Competing Interest Statement

The authors have declared no competing interest.

### Summary of Updates

This version has the correct authorship order

